# A lipid transfer-dependent feedback loop activates ATG9A compartments in autophagy initiation

**DOI:** 10.1101/2025.08.16.670665

**Authors:** Elisabeth Holzer, Justyna Sawa-Makarska, Daniel Bernklau, Julia Romanov, Martina Schuschnig, Sascha Martens

## Abstract

Autophagy degrades cellular material by sequestering it within autophagosomes, which form *de novo* from precursors called phagophores. Phagophore assembly and expansion require ATG9A-positive seed compartments, the lipid transfer protein ATG2A, and the class III phosphatidylinositol 3-phosphate kinase complex I (PI3KC3-C1). PI3KC3-C1 synthesizes phosphatidylinositol 3-phosphate (PI3P), a key lipid that drives downstream processes for phagophore expansion, including ATG8 lipidation. We find that ATG9A compartments contain only traces of phosphatidylinositol (PI), likely insufficient for efficient PI3P production or recruitment of PI3P-binding effectors. Nevertheless, ATG2A is recruited to these compartments and mediates lipid transfer, including PI, into them. Remarkably, even without detectable PI3P, ATG9A compartments are direct substrates for ATG8 lipidation, and ATG8 proteins themselves enhance ATG2A-mediated lipid transfer. In cells, ATG2A is essential for the appearance of PI3P on ATG9A compartments. Our findings support a model in which a lipid transfer-driven feedback loop activates ATG9A compartments for phagophore expansion.

## INTRODUCTION

Autophagy mediates the degradation of cellular material within lysosomes. Among the autophagic pathways, macroautophagy (hereafter autophagy) encapsulates its cargo material in double-membrane vesicles referred to as autophagosomes. Autophagosome formation can be triggered by starvation, where autophagy serves to mobilize nutrients required for cell survival. The process can also be induced by the appearance of harmful material such as protein aggregates, damaged organelles including mitochondria, and pathogens ^1^. These selective autophagy pathways ensure cellular homeostasis and prevent the development of diseases such as neurodegeneration and cancer ^2,3^.

Autophagosomes form *de novo* in proximity to the endoplasmic reticulum (ER). Initially, small, flattened membrane cisternae referred to as phagophores (also called isolation membranes) are observed ^4^. These structures gradually expand, sequestering cargo material as they grow. Upon closure, they fuse with endolysosomal compartments, where the inner membrane and enclosed cargo are degraded ^5–7^.

Autophagosome biogenesis requires the concerted action of conserved factors, collectively referred to as the autophagy machinery. The main modules of this machinery are the ULK1 kinase complex composed of the ULK1 (or ULK2) protein kinase, ATG13, ATG101, and FIP200 ^8,9^, the tetrameric class III phosphatidylinositol 3-phosphate kinase complex 1 (PI3KC3-C1) composed of the lipid kinase VPS34, VPS15, ATG14L, and Beclin1 ^10–12^, vesicles containing the ATG9A protein ^13–16^, the bridge-like lipid transfer protein ATG2A ^17–19^, the WIPI proteins including WIPI2 ^20,21^ and WIPI4 ^22^, and the ATG8 lipidation machinery including the ATG16L1 complex ^23,24^.

Among these factors, the lipid scramblase ATG9A is the only transmembrane protein ^14,25–27^. It is synthesized in the ER and trafficked to the trans-Golgi, where it is sorted into small vesicles. It cycles between the Golgi, the plasma membrane, and endosomes. Upon initiation of autophagosome formation, some of these vesicles are localized to the site of phagophore assembly to form ATG9A compartments, where they cooperate with the rest of the autophagy machinery to generate the autophagosome ^13,28–31^.

A variety of membrane sources have been implicated in autophagosome biogenesis. These include the plasma membrane ^32^, mitochondria ^33^, the Golgi apparatus ^34^ and endosomes ^35^. According to a prominent model ^16,26,27,36–39^, phagophore assembly occurs as follows ^40^. ATG9A-harboring compartments are connected to the ER via the ATG2A lipid transfer protein ^41,42^. The ULK1 and PI3KC3-C1 complexes are recruited to the ATG9A compartments, where they coordinate the activation and assembly of the rest of the machinery ^43^. In particular, the PI3KC3-C1 catalyzes the synthesis of phosphatidylinositol 3-phosphate (PI3P) from phosphatidylinositol (PI) ^44^. The PI3P in turn recruits the WIPI proteins, which stabilize the interaction of ATG2A with the membrane (in case of WIPI4) ^45,46^, or recruit the ATG16L1 complex to catalyze the lipidation of ATG8 proteins (WIPI2) ^20,24^. The growth of the initial seed membrane is driven by ATG2A-mediated transfer of lipids from the ER and the subsequent distribution of the incoming lipids across the bilayer by the ATG9A scramblase activity ^47^. This model of phagophore assembly and expansion offers an explanation for the apparent lack of transmembrane proteins in the autophagosomal membrane and the high ration of surface to lumen ^48^. In addition to ATG9A vesicles, other compartments such as cis-Golgi-derived or COPII-positive vesicles were shown to contribute to early phagophores ^34,49^.

ATG9A vesicles or compartments derived thereof have therefore a central role in phagophore assembly by acting as seed membranes to establish the initial ATG2A-dependent membrane contact site with the ER. Consistently, ATG9A compartments accumulate in large clusters together with autophagic cargoes and autophagy machinery components in ATG2A/B double knockout cells ^19,50^. In *Saccharomyces cerevisiae*, Atg9 vesicles contain high amounts of PI and can serve as substrates for the PI3KC3-C1 ^15^. The synthesized PI3P can subsequently recruit the Atg8 lipidation machinery ^51^.

Here, we set out to mechanistically dissect the role of ATG9A compartments in mammalian autophagosome biogenesis. In contrast to Atg9-positive vesicles in yeast, we find that human ATG9A compartments contain only minimal levels of PI and are not substrates for PI3KC3-C1 lipid kinase activity, even though the complex binds to them. This low PI content is particularly noteworthy, given that PI represents around 5% of total phospholipids in human cells and approximately 15% in the ER ^52^, a major membrane source for autophagosome formation. We show that ATG2A can be recruited to both ATG9A-positive compartments and reconstituted ATG9A-proteoliposomes, where it mediates lipid transfer, including PI, into these membranes. ATG9A compartments can support ATG8 lipidation despite undetectable levels of PI3P, and we find that ATG8 proteins, in turn, enhance ATG2A-mediated lipid transfer. Together, these findings support a model in which a feedback loop involving ATG2A and ATG8s activates ATG9A compartments for phagophore expansion.

Our results suggest that ATG9A compartments require activation by ATG2A-mediated lipid transfer before they become fully competent for the PI3P dependent recruitment of the autophagy machinery and subsequent phagophore expansion.

## RESULTS

### ATG9A Compartments in Mammalian Cells Contain Only Minimal PI

To investigate the nucleation of autophagosomes in mammalian cells, we aimed to reconstitute the early steps of autophagosome formation *in vitro* using purified components. In *S. cerevisiae*, autophagosome nucleation depends on Atg9-containing compartments, which can serve as platforms for the recruitment and activation of the PI3KC3-C1 ^15^. We hypothesized that a similar mechanism might operate in mammalian cells. To isolate and characterize native mammalian ATG9A compartments, we employed a CRISPR/Cas9-engineered HAP1 knock-in cell line expressing endogenously tagged ATG9A ^53^. This approach was chosen to avoid potential artifacts associated with ATG9A overexpression, which has been observed to cause mislocalization and altered trafficking ^13^. The knock-in cell line expresses ATG9A with C-terminal GFP and FLAG tags (**Fig. 1A**), preserves normal autophagic flux (**Fig. S1,A**) and enables the purification of native ATG9A compartments via FLAG-based affinity isolation (**Fig. 1A, S1B**). Negative stain transmission electron microscopy of the purified material revealed vesicles ranging from approximately 50 to 100 nm in diameter (**Fig. 1B**), consistent with the dynamic light scattering measurement (∼50 nm, **Fig. 1C**) and the expected size of endogenous ATG9A vesicles ^54^. To further characterize these structures, we performed proteomic analysis, which confirmed the enrichment of ATG9A in vesicle preparation of HAP1 ATG9A-GFP cells compared to the parental HAP1 wild-type cell line (**Fig. S1C**). Additionally, the analysis allowed to identify proteins previously reported to be associated with ATG9A compartments ^16,31,54–56^, including ATG8 proteins, the clathrin adaptor protein complexes AP1 and AP4, as well as RAB GTPases (**Fig. S1D**). To exclude contamination of ATG9A compartments with other cellular membranes, we performed western blot analysis for several membrane markers, including Calnexin (ER), GM130 (Golgi), BECLIN1 (early endosomes), TOM20 (mitochondria), TGN46 (late trans-Golgi network), LAMP1 (lysosome), and RAB7 (late endosome). These analyses showed that the FLAG-bead-bound fraction was specifically enriched for ATG9A compartments, with no detectable presence of other membrane markers (**Fig. S1E**).

**Fig. 1.**
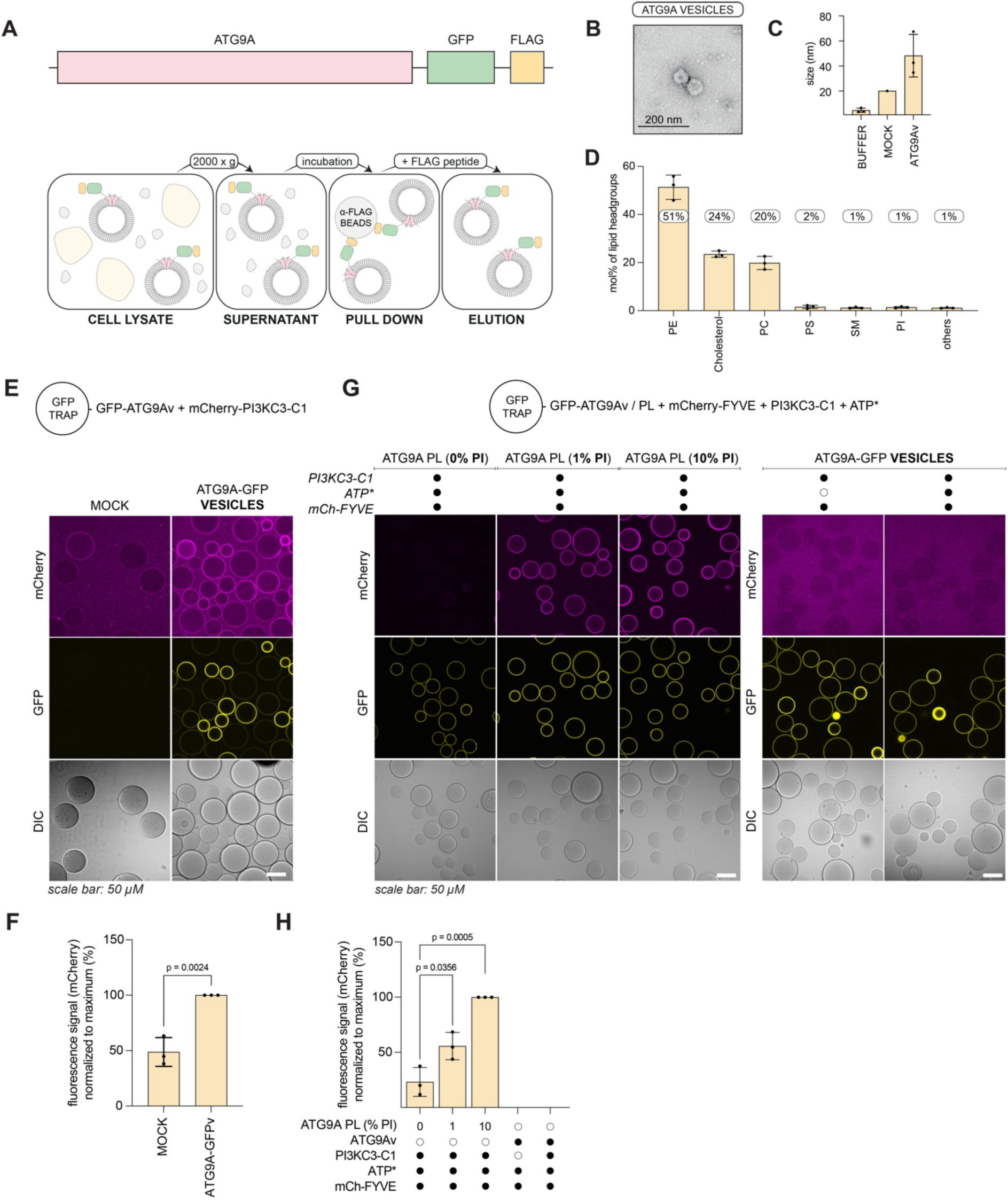
Native ATG9A compartments contain minimal levels of phosphatidylinositol. (**A**) Scheme of isolation protocol used to purify native ATG9A compartments from a HAP1 ATG9A-GFP-FLAG knock-in cell line. (**B**) Negative-stain EM image of isolated ATG9A compartments. (**C**) Dynamic light scattering analysis of ATG9A compartments compared to buffer and control sample (MOCK). Statistical significance was assessed using an unpaired *t*-test (*N* = 3). Bars represent mean ± s.d. (**D**) Quantification of lipidomic analysis of native ATG9A compartments (normalized to MOCK). To account for background, the average value from MOCK samples was subtracted of raw values (pmol) of each lipid species (*N* = 3). Background-corrected values were used to calculate molar percentage (mol%). Lipids were grouped into major classes, with additional categories including cholesteryl esters, ceramides, diacylglycerols, glucosylceramides, lactosylceramides, phosphatidylglycerol, and triacylglycerols summarized under "others". Bars represent mean ± s.d. (**E**) GFP-Trap beads coated with native ATG9A-GFP vesicles or MOCK control were incubated with mCherry-tagged PI3KC3-C1 complex (30 nM) to assess recruitment of PI3KC3-C1. **(F)** Quantification of mCherry fluorescence from panels **E** (*N* = 3) was carried out manually using ImageJ. mCherry signal (background-corrected) was expressed relative to the maximum value of each replicate. Each dot represents the mean of images acquired per replicate. Statistical significance was assessed using an unpaired *t*-test (two-tailed). Bars represent mean ± s.d. (G) GFP-Trap beads coated with ATG9A-GFP PL or native ATG9A-GFP vesicles (ATG9Av) were incubated with PI3KC3-C1 (30 nM) and mCherry-2xFYVE (30 nM) in presence or absence of ATP* (final concentrations: 0.5 mM ATP, 0.5 mM MgCl₂, 2 mM MnCl₂, 1 mM EGTA), to assess PI3KC3-C1 enzymatic activity. Recruitment of the PI3P-binding probe mCherry-2xFYVE was used as readout for PI3P production. ATG9A PLs (0/1/10% PI) were composed of 50/50/47% POPE, 20% Cholesterol, 27/26/20% DOPC, 0/1/10% Liver PI, 2% DOPS, 1% Sphingomyelin. SUVs used to prepare PL were extruded through a 100 nm filter. (H) Quantification of mCherry fluorescence from panels **G** (*N* = 3) was carried out using an AI-based image analysis pipeline. Bars represent mean ± s.d. mCherry signal (background-corrected) was expressed relative to the maximum value of each replicate. Each dot represents the mean of images acquired per replicate. Statistical significance was assessed using an unpaired *t*-test (two-tailed).

We next performed lipidomic analysis to determine the lipid composition of ATG9A-containing vesicles. In contrast to yeast Atg9 vesicles, which are enriched in (PI, ∼44%) and serve as substrates for the PI3KC3-C1 ^15^, mammalian ATG9A compartments contained only minimal amounts of PI (∼1%, **Fig. 1D**). Instead, they were primarily composed of phosphatidylethanolamine (PE, 51%), cholesterol (24%), and phosphatidylcholine (PC, 20%). To exclude the possibility that differences in the wild-type and ATG9A-GFP cell lines influenced the results, we analyzed whole cell lysates. The overall lipid profiles were highly similar, with only minor variations (**Fig. S2A, S2B**). The finding that PI remained nearly undetectable in the isolated ATG9A compartments is important, as PI3P production is required for the recruitment of downstream autophagy machinery. Our lipidomic analysis does not detect phosphorylated derivatives of PI. Thus, small amount of these lipids may be present.

Microscopy-based pull-down experiments demonstrated that the PI3KC3-C1 complex can be recruited to native ATG9A-containing vesicles (**Fig. 1E, 1F**). Recruitment of PI3KC3-C1, mediated by its multivalent membrane interactions ^57^, is independent of PI, as demonstrated by titration experiments using ATG9A-containing proteoliposomes (ATG9A PL) with 0, 1, or 10% PI, in which no significant differences in recruitment were observed (**Fig. S2C, S2D**). To assess PI3KC3-C1 activity, we monitored the recruitment of a fluorescently labeled FYVE domain ^58^, which specifically binds to PI3P (**Fig. 1G, 1H**). The absence of FYVE domain recruitment to native mammalian ATG9A compartments indicates the absence of preexisting PI3P and further that these membranes are poor substrates for PI3KC3-C1 even after its recruitment, likely due to a lack of accessible PI. Importantly, the kinase complex itself was confirmed to be functional. When added to ATG9A PLs with a lipid composition resembling that of ATG9A compartments and supplemented with 0, 1, or 10% PI, robust PI3P production was observed even the presence of only 1% PI. The mCherry-FYVE signal was stronger for 10% PI than at 1% PI, consistent with the increased PI content of the membrane (**Fig. 1G, 1H**). The ATG9A-GFP signal detected in the kinase activity assay indicated comparable amounts of ATG9A in the different samples. To further validate this observation, we performed a Western blot analysis comparing native ATG9A-containing vesicles and reconstituted ATG9A proteoliposomes. Analysis of the GFP-Trap-bound fractions revealed similar levels of ATG9A-GFP in both preparations (**Fig. S2E**). These findings suggest that the differences observed in PI3KC3-C1 activity are unlikely to be explained by substantial differences in ATG9A abundance between the samples. Furthermore, although ATG9A-positive compartments contain low levels of PI (approximately 1 mol%), this amount should in principle be sufficient to support detectable PI3P production. The lack of PI3KC3-C1 activity therefore suggests that PI may be physically inaccessible, for example due to molecular crowding or shielding within the ATG9A compartment. Such constraints may be relieved as lipid transfer proceeds and the phagophore expands.

Together, these results support the conclusion that mammalian ATG9A compartments initially lack substantial amounts of PI. It remains possible that the present PI exists in a phosphorylated form that is not a substrate for PI3KC3-C1.

### ATG2A Knockout Disrupts PI3P Localization

The low PI content in ATG9A compartments suggests that a tightly regulated lipid composition in these structures is critical for the initiation of autophagosome formation. A key candidate in this regulatory mechanism is ATG2A, a large, evolutionarily conserved lipid transfer protein that functions at membrane contact sites, particularly between the ER and nascent autophagic membranes. Although ATG2A has been well established as a lipid supplier for phagophore expansion ^17–19^, its potential role in the very early stages of this process remains less clearly defined.

To investigate whether ATG2A contributes to the formation of PI3P-enriched ATG9A-positive membranes, we analyzed the spatial relationship between autophagic membranes (LC3B-positive) and PI3P in HeLa wild-type and ATG2A/ATG2B double-knockout cells (ATG2 DKO). Consistent with previous studies ^19,50,59,60^, ATG2-deficient cells showed impaired autophagic flux, as evidenced by the accumulation LC3B-II and p62 foci (**Fig. S3A**). In these cells, large clusters of ATG9A-positive foci also accumulated, resulting from defects in phagophore expansion and autophagosome biogenesis ^16^. ATG2 DKO cells showed accumulation of several core autophagy regulators, including ATG16L1, ULK1, WIPI2, and ATG14 (**Fig. S3A**).

To determine whether PI3P generation at early autophagic membranes depends on ATG2, we analyzed the localization of the PI3P-binding probe p40PX-EGFP (**Fig. 2A**). Using the PiggyBac transposon system, HeLa wild-type and ATG2 DKO cells were stably transfected to express the probe under a doxycycline-inducible promoter, allowing fine-tuning of expression and preventing probe aggregation. To validate probe specificity, we performed co-staining with EEA1, a marker of early endosomes known to be enriched in PI3P. In both cell lines, p40PX-EGFP showed clear co-localization with EEA1, confirming its ability to detect PI3P-positive compartments (**Fig. S3B**). Under starvation conditions, LC3B-positive structures showed robust colocalization with p40PX-EGFP (Pearson correlation coefficient [PCC] ≈ 0.45) in wild-type cells, indicating that autophagic membranes are positioned near PI3P-enriched membranes and/or that they acquire PI3P (**Fig. 2B, 2C**). This observation is consistent with the recruitment of PI3P-binding effectors such as WIPI proteins, which contain WD40 domains and localize to early autophagic membranes ^61^. By contrast, colocalization between LC3B and p40PX-EGFP was markedly reduced in ATG2 DKO cell (PCC ≈ 0.18). Despite the accumulation of phagophore precursor structures, LC3B-positive vesicles in these cells exhibited minimal overlap with p40PX-EGFP, indicating a failure to acquire PI3P in the absence of ATG2 (**Fig. 2B, 2C**). To further validate the specificity of the probe and to confirm that the p40PX signal reflects PI3KC3-C1 activity, cells were treated with the VPS34 inhibitor VPS34-IN1 (2 µM, 2 h), which abolished p40PX-EGFP recruitment (**Fig. 2D**). Moreover, the defect in ATG2 DKO cells, characterized by the accumulation of phagophore precursor structures that do not co-localize with PI3P (PCC ≈ 0.21), was rescued by transient expression of wild-type ATG2A (PCC ≈ 0.46), restoring PI3P localization to LC3B-positive membranes to levels comparable to wild-type cells (PCC ≈ 0.58) (**Fig. S3C, S3D**). Additionally, we generated wild-type and ATG2 DKO cell lines co-expressing doxycycline-inducible p40PX-EGFP and mCherry-LC3B. Live-cell imaging revealed that autophagy induction led to the increase of LC3B puncta that partially co-localized with p40PX-EGFP in wild-type cells, whereas the accumulated LC3B-positive foci in ATG2 DKO cells did not. Treatment with the autophagy inhibitor VPS34-IN1 abolished p40PX-EGFP recruitment in both cell lines (**Fig. 2E**).

**Fig. 2.**
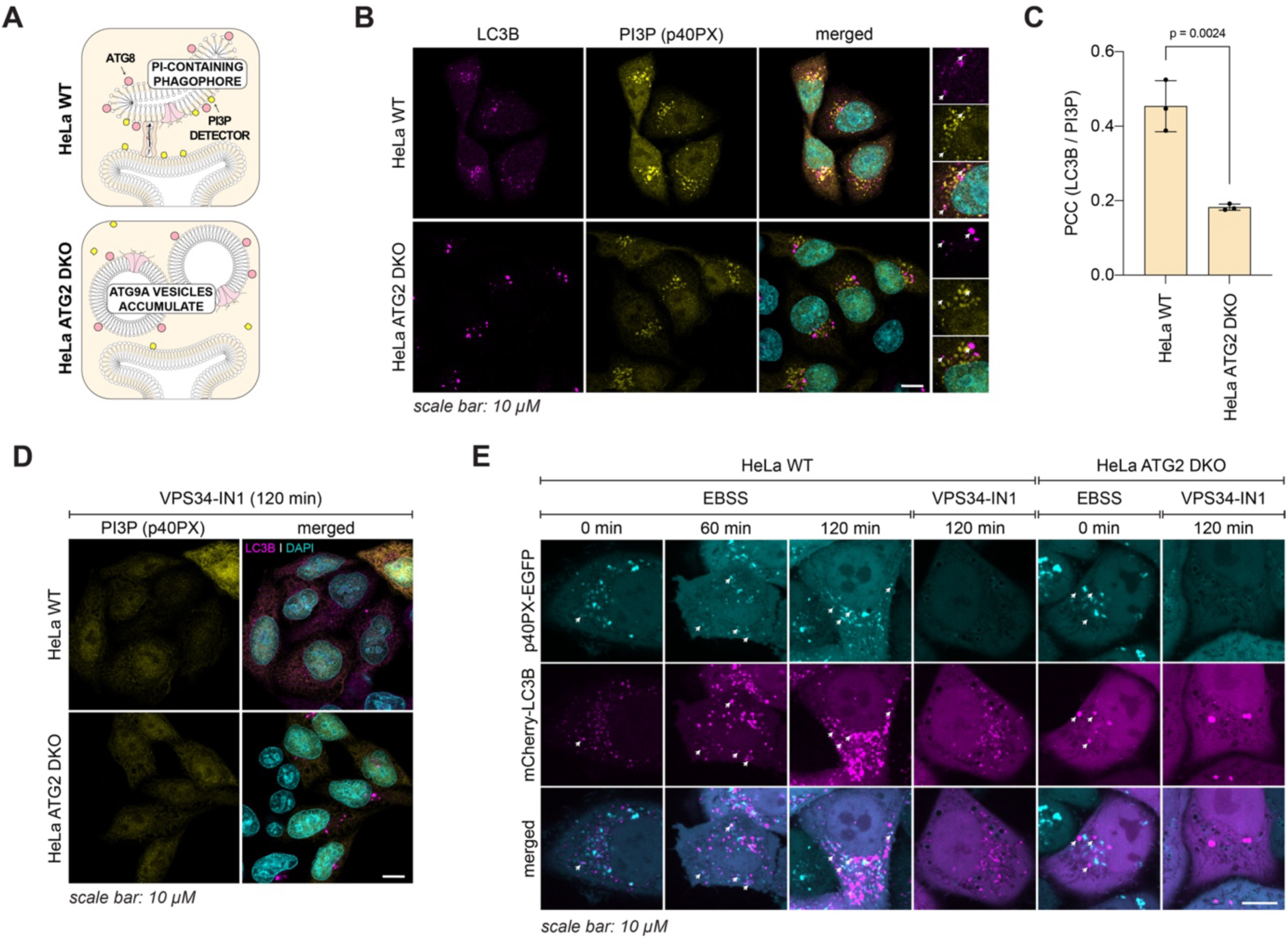
ATG2A deficiency alters PI3P localization and blocks autophagosome maturation. (**A**) Schematic model showing that in wild-type cells, PI3P-binding domains co-localize with ATG9A- and LC3B-positive compartments, consistent with PI3P enrichment. In ATG2 DKO cells, ATG9A- and LC3B-positive foci form but lack PI3P, suggesting that ATG2A is required for PI transfer into ATG9A compartments. (**B**) Immunofluorescence staining of HeLa wild-type and ATG2 DKO p40PX-EGFP (PI3P-binding domain) for analyzing the co-localization of LC3B and p40PX (**C**) Quantification (*N =* 3; 4 images per condition and replicate) of LC3B and PI3P (p40PX) co-localization shown in **B**, calculated using Pearson’s correlation coefficient (PCC). Statistical significance was assessed using an unpaired *t*-test (two-tailed). Bars represent mean ± s.d. (**D**) Immunofluorescence staining of HeLa wild-type and ATG2 DKO p40PX-EGFP cells after treatment with 2 μM VPS34-IN1 (a PI3KC3-C1 inhibitor) for 120 min. Inhibitor treatment abolishes puncta formation by p40PX, confirming PI3P-dependence. Representative images from one of three independent experiments are shown. (**E**) Live-cell images of HeLa wild-type and ATG2 DKO p40PX-EGFP mCherry-LC3B cells untreated or upon autophagy initiation by EBSS (60 or 120 min) as well as inhibition by treatment with 2 μM VPS34-IN1 (120 min). Representative images from one of three independent experiments are shown.

The clustering of WIPI2 on ATG9A-positive foci observed above (**Fig. S3A**) may therefore reflect its ability to bind not only PI3P but also other autophagy factors, such as the ULK1 kinase complex ^16,20,53,61,62^. Furthermore, the recruitment of ATG16L1 to these foci could be mediated by RAB33, which was identified in our mass spectrometry analysis of ATG9A compartments ^63^, as well as its binding to FIP200 ^64^. Notably, ATG14, a component of the PI3KC3-C1 complex, indicates that the kinase complex can be recruited to ATG9A compartments even in the absence of PI. This is consistent with our *in vitro* findings showing that PI3KC3-C1 recruitment to membranes is independent of PI (**Fig. S2C, S2D**).

Together these results show that in ATG2 DKO cells LC3B and ATG9A-positive structures (**Fig. S3A**) reside adjacent to PI3P-enriched membranes, suggesting that ATG2A facilitates PI3P accumulation on autophagosome precursors potentially by transferring PI to ATG9A-positive compartments, which is then phosphorylated by the PI3KC3-C1 complex. ATG2A may also transfer PI3P directly from the ER to phagophores during their initiation.

### ATG2A Associates with ATG9A-Positive Vesicles to Enable PI Transfer

To investigate the role of ATG2A in lipid transfer including PI into ATG9A compartments, we purified active ATG2A. The protein was N-terminally tagged with 3xFLAG and a TEV protease cleavage site (**Fig. S4A**), allowing purification using anti-FLAG affinity beads followed by on-bead cleavage with TEV protease to elute the protein (**Fig. S4B**). We confirmed that ATG2A mediates liposome tethering, which is further enhanced in the presence of WIPI4, a PI3P-binding protein and known interactor of ATG2A (**Fig. S4C**). To validate the lipid transfer activity of purified ATG2A, we employed the FRET-based lipid transfer assay, in which ATG2A-mediated lipid transfer leads to the dilution of NBD-PE into acceptor membranes (**Fig. S4D**). We also performed the assay in the presence of WIPI4, previously reported to enhance lipid transfer ^17,65^. Inclusion of WIPI4 significantly increased NBD fluorescence, confirming that the recombinant ATG2A was functionally active (**Fig. S4E**).

We next examined whether ATG2A interacts with native ATG9A compartments. ATG9A compartments were isolated from HAP1 ATG9A-GFP cells and immobilized on GFP-Trap beads. As a negative control (MOCK), HAP1 wild-type cells lacking tagged ATG9A were used. Upon incubation with recombinant ATG2A bearing a C-terminal mCherry tag, we observed robust recruitment of ATG2A to ATG9A compartments compared with the MOCK control (**Fig. 3A, 3B**), presumably at least in part via ATG9A ^41,42^. We then employed DOPE lipids labeled with a second FRET pair, ATTO540Q and ATTO425, which can be spectrally separated from the GFP signal of ATG9A, to monitor lipid transfer into native ATG9A compartments. In donor SUVs, ATTO425 fluorescence is quenched by ATTO540Q. Upon lipid transfer from donor vesicles to ATG9A compartments, dilution of the fluorophores leads to an increase in ATTO425 fluorescence (**Fig. 3C**). We validated the new FRET pair in a standard lipid transfer assay in which donor SUVs were mixed with acceptor SUVs in the absence or presence of ATG2A (**Fig. 3D**). Addition of ATG2A resulted in increased ATTO425 fluorescence. Notably, although ATG9A compartments alone were able to recruit donor SUVs in the absence of ATG2A, lipid transfer was strongly enhanced in the presence of ATG2A, as indicated by the increased ATTO425 signal (**Fig. 3E, 3F**). Together, these results demonstrate that isolated ATG9A compartments recruit ATG2A and are functionally competent for lipid transfer.

**Fig. 3.**
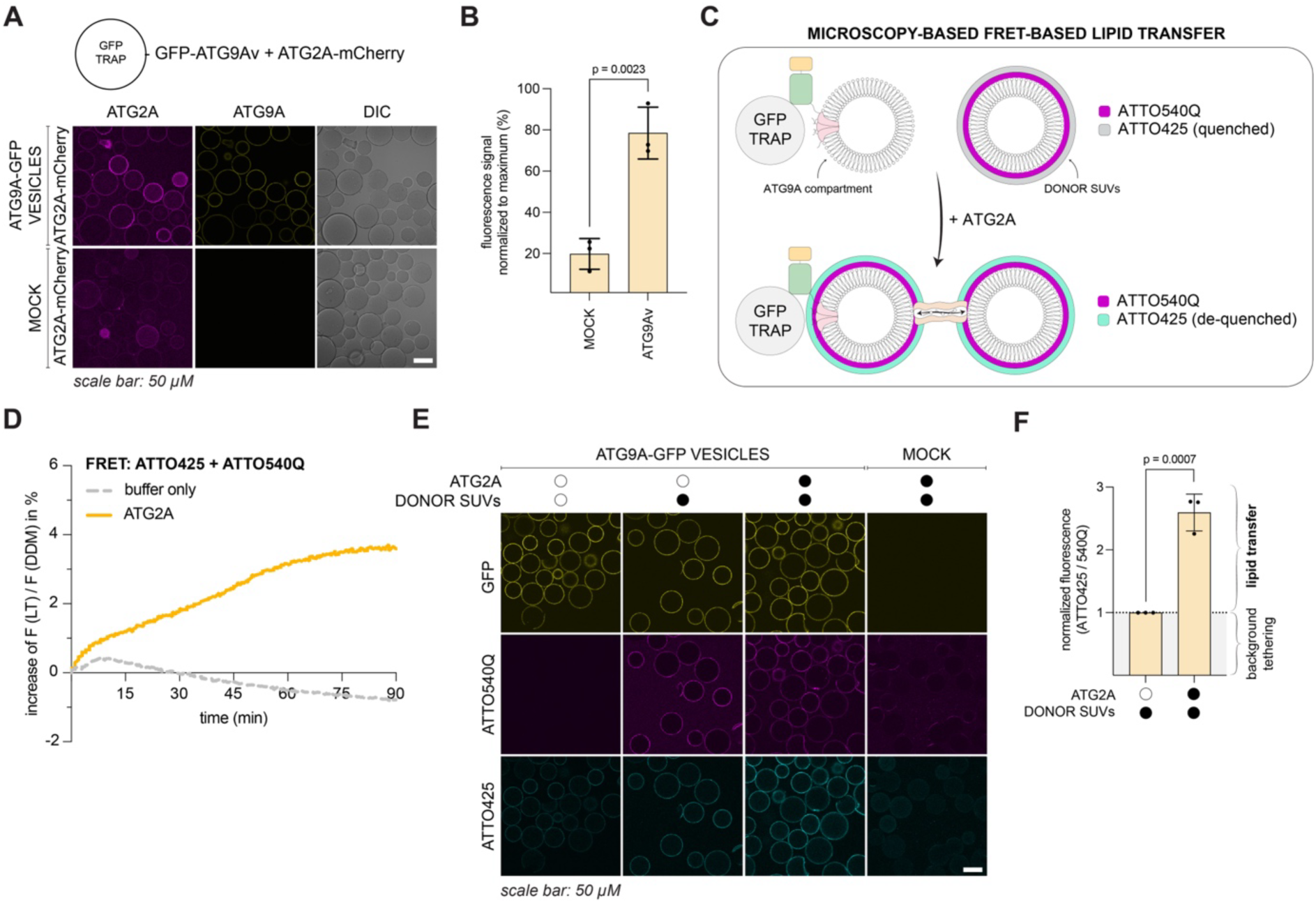
ATG2A tethers membranes and mediates lipid transfer. (**A**) GFP-Trap beads coated with native ATG9A-GFP vesicles or MOCK control were incubated with mCherry-ATG2A (30 nM) to assess recruitment of ATG2A. (**B**) Quantification of mCherry fluorescence from ATG2A recruitment in **A** (*N =* 3; independent experiments) was carried out manually using ImageJ. Total fluorescence signals were background-corrected and normalized within each replicate to the respective maximum value. Statistical significance was assessed using an unpaired *t*-test (two-tailed). Bars represent mean ± s.d. (**C**) Schematic of FRET-based lipid transfer to ATG9A compartments. Isolated ATG9A-GFP compartments were immobilized to GFP-trap beads and incubated with donor SUVs in the presence or absence of ATG2A. The donor SUVs contained an ATTO425/ATTO540Q FRET pair (66% DOPC, 30% DOPE, 2% ATTO425-DOPE, 2% ATTO540Q-DOPE; 100 nm in size), in which ATTO425 is initially quenched by ATTO540Q. (**D**) Lipid transfer activity of ATG2A was monitored at 470 nm (excitation at 405 nm) and normalized to the fluorescence after membrane solubilization with 0.5% DDM (F(DDM)). Acceptor SUVs contained 65% DOPC, 15% DOPS, 15% DOPE, and 5% PI3P, whereas donor SUVs contained 66% DOPC, 30% DOPE, 2% ATTO425-DOPE, and 2% ATTO540Q-DOPE (100 nm in size). Statistical significance was assessed using an unpaired *t*-test (*N* = 3). Bars represent mean ± s.d. (**E**) Microscopy images of GFP-Trap agarose beads coated with isolated ATG9A compartments that were incubated with donor SUVs containing an ATTO425/ATTO540Q FRET pair in presence or absence of ATG2A (as shown in **C**). The condition ATG9A-compartmens-alone is used to assess bleed through of GFP containing ATG9A compartments. (**F**) Quantification (*N =* 3; independent experiments) of lipid transfer shown in **E**. Intensities in red (ATTO540Q) and blue (ATTO425) channels were quantified to monitor SUV tethering and lipid transfer. ATTO425 signal was corrected for bleed-through using ATG9A compartments-only control. Lipid transfer was assessed as ATTO425/ATTO540Q ratio, thereby normalizing transfer to the amount of bound donor SUVs. The resulting values were normalized to no-ATG2A condition, where residual blue signal arises from incompletely quenched ATTO425 in donor SUVs. Values above 1 indicate dequenching of ATTO425, resulting from ATG2A-mediated lipid transfer. Statistical significance was assessed using an unpaired *t*-test (two-tailed). Bars represent mean ± s.d.

To determine whether ATG2A can mediate the transfer of PI as a substrate for PI3KC3-C1 into ATG9A-containing membranes, we performed a stepwise reconstitution assay (**Fig. 4A**). Donor SUVs containing PI and ATTO390-PE were first incubated with ATG2A and ATG9A PL to initiate lipid transfer (step 1). After immobilization of ATG9A PL on GFP-Trap beads, donor SUVs and ATG2A were removed by several washing steps. The remaining ATG9A PL were incubated with the PI3KC3-C1 complex, cofactors (ATP*) and the PI3P-binding probe mCherry-FYVE to detect synthesized PI3P on ATG9A PL (step 2). A robust ATTO390 signal was detected only in samples that included ATG2A during the initial incubation step, indicating lipid transfer. Importantly, robust recruitment of the FYVE domain was observed only when all components – donor SUVs, ATG2A, PI3KC3-C1 and ATP* – were present, consistent with PI delivery to ATG9A membranes and subsequent phosphorylation by PI3KC3-C1 (**Fig. 4B, 4C**). Two independent experiments strongly suggest that the observed signal predominantly reflects lipid transfer into ATG9A membranes. First, the amount of ATG2A-mCherry associated with ATG9A PL was strongly reduced after the washing steps, indicating efficient removal of tethered donor SUVs (**Fig. S5A**). Second, bulk lipid transfer into ATG9A PL was independently confirmed using the FRET-based assay with ATTO540Q and ATTO425 (**Fig. S5B**). Upon addition of ATG2A, we observed an increase in ATTO425 fluorescence, indicating lipid transfer between donor SUVs and ATG9A PL (**Fig. S5C, S5D**). We also attempted to separate the lipid transfer and membrane tethering functions of ATG2A by testing previously published lipid transfer mutants ^42,66,67^. However, while these mutants displayed strongly impaired lipid transfer activity *in vitro*, they also exhibited defects in membrane tethering in our assays, preventing a direct assessment of lipid transfer independent of tethering.

**Fig. 4.**
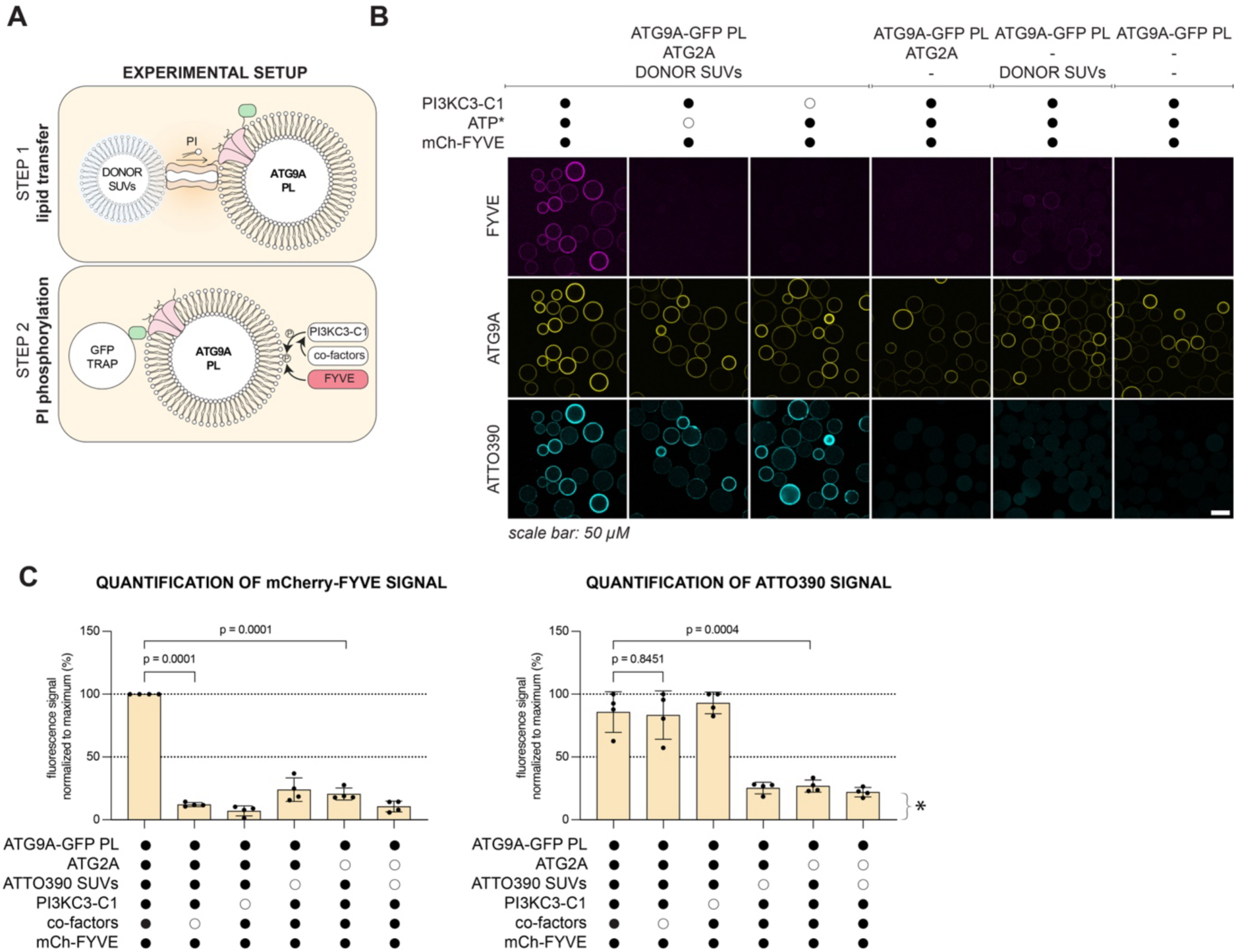
ATG2A transfers PI-containing lipids into reconstituted ATG9A compartments and facilitates PI3P production. (**A**) Schematic of the experimental setup shown in **B**. (**B**) ATG9A PL were incubated for 2.5 hours with ATG2A and donor SUVs containing PI and ATTO390-PE. Control conditions included: ATG2A only, donor SUVs only, or buffer alone. After incubation, the proteoliposomes were immobilized on GFP-Trap beads (2.5 h), washed, and incubated with 30 nM PI3KC3-C1, 30 nM mCherry–FYVE, and cofactors (ATP*) to assess PI transfer and PI3P generation. ATG9A PLs were composed of 50% POPE, 20% Cholesterol, 27% DOPC, 0% Liver PI, 2% DOPS, 1% Sphingomyelin. This composition was designed to mimic native ATG9A compartments, while being devoid of PI, thereby enabling the assessment of PI transfer. ATTO390- and PI-containing donor SUVs were composed of 56.5% DOPC, 16% DOPE, 16% DOPS, 10% liver PI, and 1.5% ATTO390-PE. (**C**) Quantification (*N =* 4; independent experiments) of mCherry fluorescence (left panel; PI3P detection via FYVE binding) and ATTO390 fluorescence (right panel; lipid transfer) from **B**. Statistical analysis was performed using an unpaired *t*-test (two-tailed). Bars represent mean ± s.d. Quantification was carried out using an AI-based image analysis pipeline. The asterisk in the quantification of the ATTO390 signal indicates technical background due to bleed-through from the 488 nm into the 405 nm channel.

To summarize, these results support a role for ATG2A-mediated PI transfer to ATG9A compartments, rendering them competent for downstream PI3KC3-C1-mediated PI phosphorylation.

### PI3P Enrichment on LC3B-Positive Autophagic Membranes Requires ATG2A Function

The minimal levels of PI and PI3P on ATG9A compartments during the early stages of phagophore nucleation suggest that WIPI4, which requires PI3P for membrane binding, is not involved in the initial recruitment or stabilization of ATG2A. Supporting this, when cells were transfected with EGFP-WIPI4, we did not observe clustering of WIPI4 with ATG9A-positive foci in HeLa ATG2 DKO cells (**Fig. S6A**). Furthermore, although ATG2A and WIPI4 have been reported to form a complex ^46^, our pull-down assay (**Fig. S6B**) and analytical size-exclusion chromatography (**Fig. S6C, S6D**) indicate that this interaction is relatively weak. These findings suggest that, in mammalian cells, additional mechanisms are likely required for ATG2A recruitment. ATG8 proteins are candidates for this activity as ATG2A contains a functional LC3-interacting region (LIR) adjacent to its WIPI4-binding site ^68^, suggesting that ATG8 proteins may facilitate the early association of ATG2A with membranes.

Because ATG2A is required for bulk lipid transfer during phagophore expansion, ATG9A-positive compartments accumulating in ATG2 DKO cells are expected to represent early autophagy precursors that have not yet received substantial lipid input. We therefore reasoned that these compartments may resemble the initial substrate for ATG8 lipidation.

To investigate the nature of ATG9A-positive compartments in ATG2-deficient cells, we first characterized the isolated vesicles using a protease protection assay and electron microscopy. For efficient isolation of ATG9A-positive compartments, we used HAP1 ATG9A-GFP and HAP1 ATG9A-GFP ATG2 DKO starved cells in which we blocked late steps of autophagy by treating them with Bafilomycin A. Protease sensitivity of any cargo was assessed using the autophagy cargo p62 as a marker. In fractions derived from HAP1 ATG9A-GFP cells, p62 was resistant to proteinase K digestion, indicating sequestration within sealed autophagosomes. In contrast, consistent with defect in phagophore expansion ^16^, p62 remained protease-sensitive in fractions from ATG2 DKO cells (**Fig. 5A, 5B**). To determine whether the loss of ATG2 affects the gross morphology of the isolated ATG9A compartments, we performed electron microscopy. ATG9A-positive vesicles from wild-type and ATG2 DKO cells appeared morphologically similar, supporting the idea that ATG2 deficiency primarily affects membrane elongation rather than the formation of the initial vesicular structures (**Fig. 5C**).

**Fig. 5.**
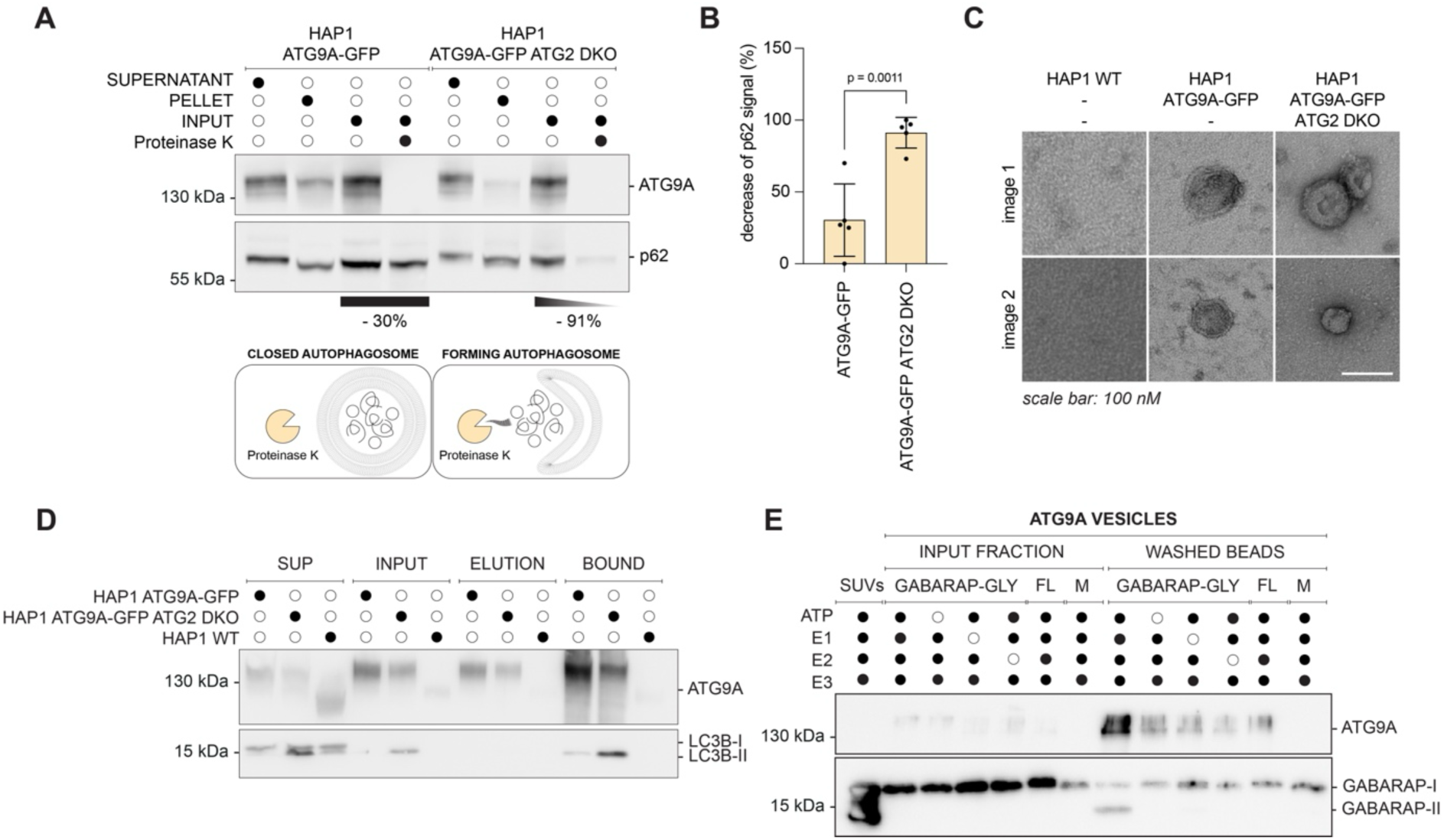
Loss of ATG2 disrupts autophagosome closure and PI3P enrichment. (A) Western blot analysis of isolated autophagosomes from HAP1 ATG9A-GFP and HAP1 ATG9A-GFP ΔATG2 cells before and after Proteinase K (PK) digestion. In ATG9A-GFP cells, approximately 30% of the p62 signal is lost, indicating the presence of closed, protease-protected vesicles. In contrast, ATG2 DKO cells show ∼91% loss of p62. SUPERNATANT refers to the soluble fraction collected during the final isolation step, whereas PELLET refers to the unwashed AP fraction. INPUT corresponds to the washed APs used for the PK digest. (B) Quantification of p62 signal following PK digestion, as shown in **B** (*N =* 5; independent experiments). Statistical significance was determined using an unpaired *t*-test (two-tailed). Bars represent mean ± s.d. (**C**) Negative-stain EM image of isolated ATG9A compartments from HAP1 ATG9A-GFP and HAP1 ATG9A-GFP ATG2 DKO cells. Representative images from one of two independent experiments are shown. (**D**) Western blot analysis of samples collected during isolation of native ATG9A compartments. Shown are supernatant (SUP), input fraction for FLAG-Trap incubation (INPUT), FLAG-Trap elution (ELUTION), and final vesicle fraction bound to GFP-Trap beads (BOUND) from HAP1 WT, HAP1 ATG9A-GFP, and HAP1 ATG9A-GFP ATG2 DKO cells. ATG9A is enriched in the BOUND fraction of both ATG9A-GFP and ATG2 DKO cells compared to WT. LC3B signal is increased in the ATG9A-containing vesicles from ATG2 DKO cells, suggesting the accumulation of LC3-positive, unsealed membranes. Representative images from one of three independent experiments are shown. (**E**) Western blot analysis of GABARAP conjugation to ATG9A vesicles. GABARAP was chosen as the substrate because its lipidated form (GABARAP-II) exhibits a characteristic mobility shift relative to the unlipidated form (GABARAP-I). GFP-Trap agarose beads coated with ATG9A vesicles were incubated with GABARAP, ATG7 (E1), ATG3 (E2), ATG12–5–16L1 complex (E3), and ATP. The lipidation reaction was controlled by individually omitting each essential component of the lipidation machinery (E1, E2, or E3). PE-containing SUVs served as a positive control for lipidation. GABARAP-GLY: GABARAPΔL117 mutant exposing the penultimate glycine for conjugation; FL: full-length GABARAP, which cannot undergo lipidation; M: mock-purified control from HAP1 wild-type cells. Representative images from one of three independent experiments are shown.

We next examined whether ATG8 proteins are present on ATG9A compartments. Isolation of these structures from HAP1 ATG9A-GFP ATG2 DKO cells revealed a strong enrichment of lipidated LC3B compared to HAP1 ATG9A-GFP cells (**Fig. 5D**).

Consistent with this, Olivas and colleagues used nanodiscs to demonstrate that ATG9A and LC3B-II can reside on the same membrane even in ATG2 DKO cells. Their findings indicate that LC3B lipidation can occur on ATG9A-positive membranes before ATG2-dependent membrane expansion^16^. While that study did not directly address the PI3P status of these structures, it supports the notion that ATG9A-containing membranes can serve as substrates for LC3 lipidation at an early stage of autophagosome biogenesis.

To directly test if ATG9A compartments can serve as substrates for ATG8 lipidation, we performed *in vitro* conjugation assays using ATG7 (E1), ATG3 (E2), and the ATG12–5– 16L1 complex (E3). As a substrate, we used GABARAPΔL117, which exposes the penultimate glycine and exhibits a distinct gel shift upon lipidation. ATG9A compartments isolated from HAP1 ATG9A-GFP cells under nutrient-rich conditions supported robust GABARAP lipidation (**Fig. 5E, S6E, S6F**), indicating that they provide a suitable lipid environment, likely due to their high PE content (**Fig. 1D**).

### ATG8 Directly Interacts with ATG2A and Enhances Its Lipid Transfer Activity

Having established that ATG8 proteins are present on ATG9A-positive membranes prior to PI3P enrichment, we investigated whether they directly contribute to ATG2A recruitment and/or activation. Using *in vitro* binding assays, we compared wild-type ATG2A with a mutant lacking a functional LC3-interacting region (LIR) motif, in which amino acids 1362–1365 were mutated to alanines ^68^. Wild-type ATG2A bound strongly to multiple ATG8 family members, whereas the LIR mutant showed markedly reduced interactions, particularly with LC3A and LC3B, while binding to GABARAP and GABARAPL1 was still preserved (**Fig. 6A, 6B, S7A**). These results suggest that ATG2A engages ATG8 proteins via different binding motifs.

**Fig. 6.**
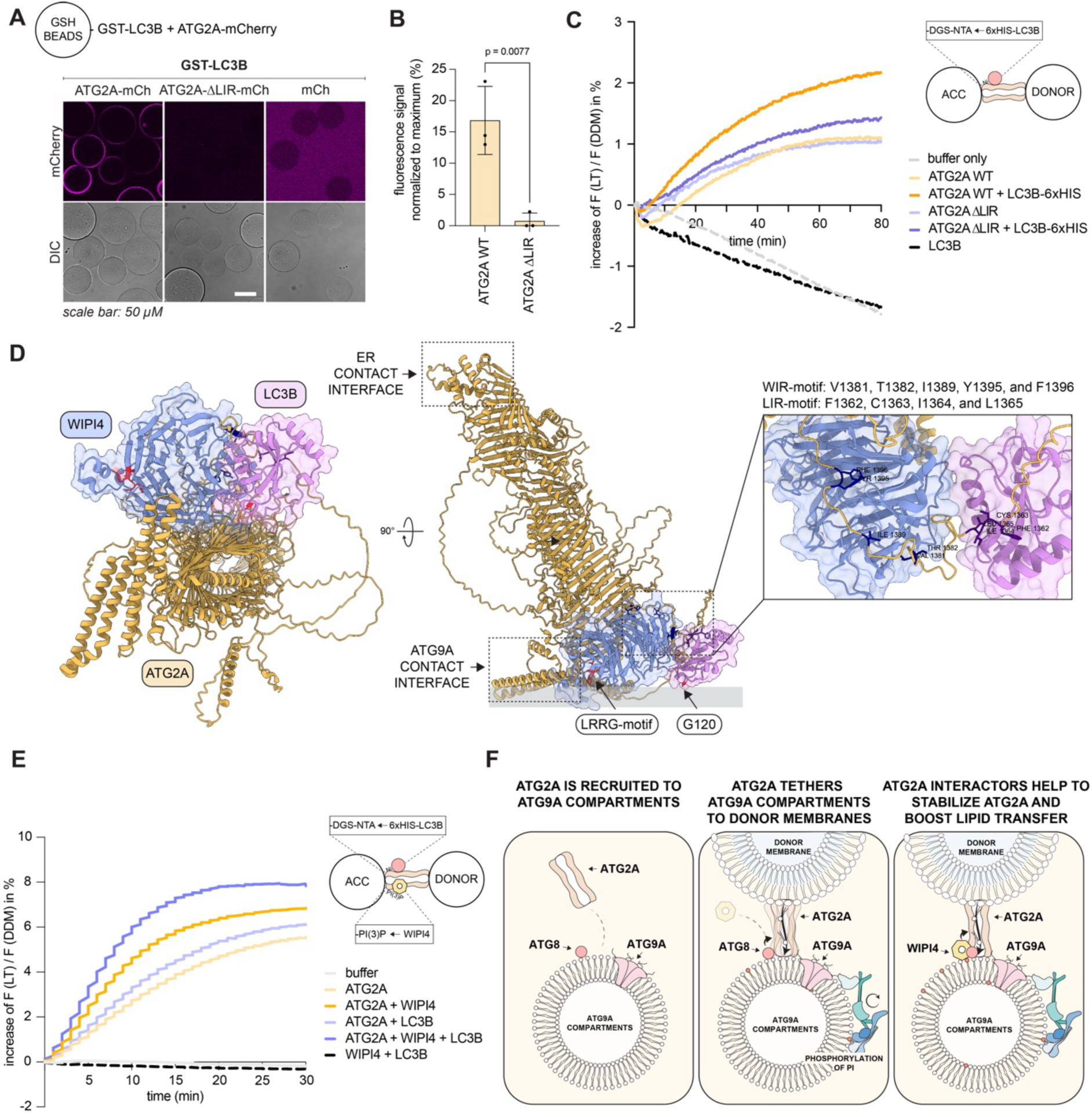
ATG8 directly interacts with ATG2A and enhances its lipid transfer activity. (**A**) GSH-Trap beads coated with GST-tagged LC3B were incubated with ATG2A-mCherry, ATG2A-ΔLIR-mCherry (aa 1362–1365; FCIL/AAAA) or mCherry. (**B**) Quantification of mCherry fluorescence signal from ATG2A recruitment in **A** (*N =* 3; independent experiments). Quantification was carried out using an AI-based image analysis pipeline. Total fluorescence signals were normalized within each replicate to the respective maximum value. Each dot represents the mean of images acquired per replicate. Statistical significance was assessed using an unpaired *t*-test (two-tailed). Bars represent mean ± s.d. (**C**) Comparison of ATG2A WT and ΔLIR lipid transfer activity in absence or presence of 6XHIS-LC3B (100 nM). Acceptor SUVs contain 60% DOPC, 15% DOPS, 15% DOPE, 5% PI3P and 5% DGS-NTA and donor SUVs contained 66% DOPC, 30% DOPE, 2% NBD-DOPE, and 2% Rhodamine B-DOPE (100 nm in size). NBD fluorescence (F(LT)) was recorded at 535 nm (excitation at 485 nm) and normalized to DDM-solubilized maximum fluorescence (F(DDM)). Lines represent mean values (N = 4; independent experiments). (D) AlphaFold3-predicted structure of the ATG2A-WIPI4-LC3B complex 45,68. The model has a predicted inter-chain TM-score (ipTM) of 0.79 and TM-score (pTM) of 0.56. The predicted alignment error (PAE) heatmap is shown in Fig. S7C. (E) Comparison of lipid transfer activity between ATG2A (250 nM) in the absence or presence of WIPI4 (100 nM), 6xHIS-LC3B (100 nM) as well as WIPI4 or 6xHIS-LC3B alone (100 nM). NBD fluorescence (F(LT)) was recorded at 535 nm (excitation at 485 nm) and normalized to fluorescence after membrane solubilization with 0.5% DDM (F(DDM)). Acceptor and donor SUVs had the same composition as described in (C). Lines represent mean values (N = 4; independent experiments). (F) Schematic model of ATG2A-mediated lipid transfer and autophagosome growth. ATG2A is recruited to early autophagic membranes through interactions with ATG9A compartments and membrane-anchored ATG8 proteins. Once localized, ATG2A transfers PI or PI3P from donor membranes into ATG9A-positive compartments. Locally synthesized PI3P recruits WIPI4, which in turn helps to stabilize the membrane association of ATG2A. This positive feedback loop enhances lipid transfer activity, promoting membrane expansion and facilitating the growth of the nascent autophagosome.

To determine whether the interaction between ATG2A and ATG8 proteins modulates lipid transfer, we performed FRET-based lipid transfer assays. The addition of LC3B significantly enhanced lipid transfer by wild-type ATG2A, an effect that was almost abolished for the ΔLIR mutant (**Fig. 6C**). This enhancement likely resulted from improved membrane tethering, as supported by tethering assays (**Fig. S7B**), and suggests that LC3B binding promotes ATG2A-mediated lipid transport.

To investigate whether WIPI4 and LC3B can bind ATG2A simultaneously, we performed AlphaFold3 modeling using full-length ATG2A in complex with both proteins (**Fig. 6D, S7C**). The resulting structure predicted a stable ternary complex in which LC3B binds to the LIR (aa 1362–1365) motif of ATG2A ^68^, while WIPI4 engages the WIPI-interacting region (WIR) near the C-terminus ^45^. The model had a predicted inter-chain TM-score (ipTM) of 0.79 and a pTM of 0.56, with no steric clashes, indicating a feasible and cooperative binding configuration. Notably, LC3B and WIPI4 were positioned adjacently along a C-terminal loop region of ATG2A, supporting the idea that they can simultaneously cooperate in ATG2A recruitment or regulation of lipid transfer. This notion is further supported by the FRET-based lipid transfer assay, which showed a pronounced increase in lipid transfer when both factors were present compared with conditions in which only one interactor was present (**Fig. 6E**).

Together, these findings support a model in which ATG8 proteins localize to ATG9A-containing precursor membranes during the early stages of autophagosome formation, where they contribute to the recruitment and activation of ATG2A. By directly binding ATG2A, ATG8 proteins enhance the lipid transfer by ATG2A, facilitating the delivery of PI to nascent membranes. PI is subsequently phosphorylated to PI3P, enabling the recruitment of WIPI4. WIPI4 further recruits ATG2A, establishing a positive feedback loop that amplifies lipid transfer. As PI3P levels rise, they promote additional ATG8 lipidation and drive phagophore expansion, thereby accelerating autophagosome biogenesis (**Fig. 6F**).

## DISCUSSION

Autophagosome biogenesis requires the complex interplay of various components of the autophagy machinery. Among the core autophagy factors, ATG9A is the only conserved transmembrane protein essential for autophagosome formation. ATG9A compartments are distributed throughout mammalian cells and participate in various processes. A substantial body of evidence points to ATG9A compartments as nucleation sites for phagophore assembly ^15,16^. ATG9A is recruited early during phagophore nucleation, and in yeast Atg9 localizes to ER-proximal sites of the growing phagophore ^16,36–39,69^. Additionally, ATG9A binds to the C-terminal region of ATG2A, while the N-terminus of ATG2A is thought to associate with the ER^41,42,70^.

Our finding that ATG9A compartments are poor in PI for PI3P generation suggests that they are refractory to spurious PI3P production and subsequent recruitment of the autophagy machinery. Upon clustering and activation of the autophagy machinery either during starvation or selective autophagy at cargo sites, the network of interactions involving the ULK1 complex may establish transient membrane contact sites between ATG9A compartments and the ER via ATG2A ^65,71,72^. These contact sites likely involve binding of the ATG2A C-terminal region to ATG9A ^41,42^ and may also depend, at least in part, on the interaction of ATG2A with ATG8 proteins that are present on these compartments.

The LIR motif mediating the ATG8 interaction site resides in a loop within the C-terminus of ATG2A ^68^, consistent with ATG2A engaging both ATG9A and ATG8s on the same membrane. We detect some levels of ATG8 in ATG9A compartments in the absence of an autophagy stimulus (**Fig. S1D, 5D**). This could reflect either a constitutive feature of these vesicles or indicate that a subset of ATG9A compartments is involved in basal autophagy. In ATG2A/B-depleted cells, autophagy machinery components including WIPI2 and the ATG16L1 complex (**Fig. S3A**), localize to ATG9A-containing foci, likely representing abortive autophagosome biogenesis sites. The recruitment of these proteins could occur through multiple PI3P-independent interactions, including binding of WIPI2 to the ULK1 protein kinase complex ^16,20,53,61,62^ or association of ATG16L1 with RAB33 ^63^ and FIP200 ^64^.

Consistent with previous findings ^16^, our data show that ATG9A compartments isolated from cells lacking ATG2A display markedly increased levels of lipidated LC3B (**Fig. 5D**), suggesting that, in the absence of ATG2A-mediated lipid transfer, these compartments can act as substrates for ATG8 lipidation. Supporting this notion, we show that ATG9A compartments contain PE and PS (**Fig. 1D**), both substrates for lipidation reactions, and can serve as direct targets for ATG8 lipidation *in vitro* (**Fig. 5E, S6E, S6F**). In contrast, previous biochemical work in yeast demonstrated that Atg9 vesicles alone are not sufficient to support Atg8 lipidation and instead require PE transfer by Atg2 ^15^. Moreover, it has been shown that liposomes containing 50% PE are efficient substrates for the lipidation machinery *in vitro*, even in the absence of the ATG16L1 complex ^73^. Mammalian ATG9A compartments may have a more favorable basal capacity to support ATG8 conjugation compared with their yeast counterparts.

It is therefore possible that local clustering and activation of the ATG16L1 complex stimulates ATG8 attachment to ATG9A compartments, which in turn stabilizes the association of ATG2A. This weak network of interactions may ensure that lipid transfer from the ER only occurs at sites where the autophagy machinery remains clustered long enough. Lipid transfer from the ER may also provide PI as a substrate for the PI3KC3-C1 complex, which is recruited and activated at the ATG9A compartment via the ULK1 complex. Consistently, it has previously been shown that phosphatidylinositol synthase (PIS) is enriched at sites near autophagosome biogenesis ^74^. The PI3P generated may recruit WIPI4, which, through a positive feedback loop, stabilizes the interaction between ATG2A and the ATG9A compartment and markedly enhances the lipid transfer activity of ATG2A during phagophore expansion. Notably, the ATG2A-WIPI4 interaction is relatively weak (**Fig. S6B-D**). Thus, WIPI4 may be recruited downstream of PI transfer. In yeast, the WIPI4 homolog Atg18 has been shown to be recruited after initial membrane contact site formation mediated by Atg2 and TRAPPIII complex ^75^. The weak ATG2A-WIPI4 interaction may additionally allow ATG2A to function in various other processes where WIPI4 is not required.

Furthermore, PI3P on the phagophore can enhance the recruitment of WIPI2, which promotes PI3KC3-C1 activity and binds ATG16L1 to facilitate ATG8 lipidation, thereby further stabilizing the lipid transfer complex and enhancing recruitment of the autophagy machinery ^20,21^. An additional feedback loop may be driven by the transfer of disordered lipids from the ER into more tightly packed, post-Golgi-derived ATG9A compartments. This could facilitate the recruitment of autophagy factors such as PI3KC3-C1, WIPIs, ATG16L1, and ATG3, all of which rely on membrane insertion for their association with membranes.

Our model proposes that PI is transported from the ER into ATG9A compartments to support PI3P synthesis on the phagophore. While PI3P is clearly localized to autophagosomal membranes ^76–78^, we cannot exclude the possibility that, particularly during the early stages of phagophore nucleation, PI3P is generated at the ER and subsequently transported to ATG9A compartments via ATG2A. Activation of PI3KC3-C1 requires membrane-localized RAB1, which is present on the ER ^57,79^. Consistently, we detect RAB1 in our proteomics dataset of isolated ATG9A compartments (**Fig. S1D**), potentially enabling direct engagement of PI3KC3-C1 with these membranes. PI3KC3-C1 activity is also regulated by reversible post-translational modifications, including phosphorylation and lactylation ^80–83^. These regulatory mechanisms may represent additional layers of control influencing PI3P production. Their potential contribution to the observed phenotypes warrants further investigation in future work. Moreover, it remains possible that ATG9A compartments undergo a prefusion event with a second compartment that delivers PI such as cis-Golgi or ERGIC-derived structures ^34,49^, thereby rendering them competent for PI3P synthesis. However, the bulk amount of PI and/or PI3P is likely derived from the ER because we show here that the ATG9A-postive clusters in ATG2 DKO cells are negative for PI3P. These models above are not mutually exclusive, and future research will likely reveal additional feedback loops ensuring the precise spatial and temporal regulation of autophagosome biogenesis.

## Methods

### Plasmid Construction

Coding sequences for all constructs were amplified by PCR from existing plasmids. For expression in insect cells, codon-optimized versions were commercially synthesized (GenScript). Plasmid construction was carried out using Gibson assembly. Inserts and vector backbones were generated either by PCR or by excision from agarose gels following restriction digestion (1 h at 37 °C). Purification of DNA fragments was performed using the Wizard® SV Gel and PCR Clean-Up System (Promega, Cat# A9282). Equimolar amounts of purified insert and backbone were assembled using the NEBuilder® HiFi DNA Assembly Master Mix (New England Biolabs, Cat# E2621S) according to the manufacturer’s protocol. The resulting Gibson reactions were incubated for 1 h at 50 °C and transformed into *E. coli* DH5α competent cells (Thermo Fisher Scientific, Cat# 18265017).

Transformed cells were plated on LB-agar containing the appropriate antibiotic (ampicillin, kanamycin, or chloramphenicol) and incubated overnight at 37 °C. Individual colonies were picked and cultured overnight in LB medium. Plasmid DNA was extracted using the GeneJet Plasmid Miniprep kit (Thermo Fisher, Cat# K0503). All inserts were confirmed by Sanger sequencing (Microsynth AG), and selected clones were further validated by whole plasmid sequencing (Microsynth AG). A detailed protocol is available on protocols.io (dx.doi.org/10.17504/protocols.io.8epv5x11ng1b/v1).

### Cell lines

Cells were maintained at 37 °C in a humidified atmosphere with 5% CO₂. HAP1 cells (RRID: CVCL_Y019) were received from Horizon Discovery and HeLa cells (RRID: CVCL_0058) were obtained from the American Type Culture Collection (ATCC). The HeLa ATG9A KO cell line was a kind gift from Michael Lazarou (CVCL_F0LG). HAP1 cell lines were cultured in Iscove’s Modified Dulbecco’s Medium (IMDM; Thermo Fisher, Cat# 31980048), supplemented with 10% (v/v) fetal bovine serum (FBS; Sigma-Aldrich, Cat# F7524) and 1% (v/v) penicillin– streptomycin (Sigma-Aldrich, Cat# P4333). HeLa cell lines were grown in Dulbecco’s modified Eagle medium (DMEM, Thermo Fisher, Cat# 41966029) supplemented with 10% (vol/vol) FBS, 25 mM HEPES (Thermo Fisher, Cat# 15630080), 1% (vol/vol) non-essential amino acids (Thermo Fisher, Cat# 11140050) and 1% (vol/vol) penicillin–streptomycin. All cell lines were routinely tested for mycoplasma contamination. A detailed protocol is available on protocols.io (dx.doi.org/10.17504/protocols.io.n2bvj3y5blk5/v1).

### Generation of HAP1 ATG9A-GFP cell line

The genomic DNA of HAP1 wild-type cells was tagged at the ATG9A C-terminus locus (RRID: CVCL_E2TR) using CRISPR/Cas9 technology, providing a homology repair template containing the desired tag. A plasmid-based homology repair template (Addgene_244942) and the AIO-Puro Cas9 Nickase plasmid (Addgene_74630, ^84^) including the selected guide RNAs (gRNA-A CCTCTTCGGGCACGGGCTCA, gRNA-B: GGAGGATGAGCTACCCCCTC) were transfected to HAP1 cells. Cells were selected for Puromycin resistance (Thermo Fisher, Cat# A1113803), FACS sorted for GFP expression and best clones selected based on gDNA analysis and WB characterization. A detailed protocol is available on protocols.io (dx.doi.org/10.17504/protocols.io.3byl46j4ogo5/v1).

### Generation of CRISPR/Cas9 ATG2A/ATG2B knockout cell lines

HAP1 ATG9A-GFP ATG2A/ATG2B double knockout clone #1.3 (CVCL_F0LE) and HeLa ATG2A/ATG2B double knockout clone #7 (CVCL_F0LF) were generated with CRISPR/Cas9. Candidate single-guide RNAs (sgRNAs) were identified using CRISPick (RRID:SCR_025148; https://portals.broadinstitute.org/gppx/crispick/public), targeting all common splicing variants. The sgRNAs were ordered as short oligonucleotides (Microsynth AG) and cloned into pSpCas9(BB)-2A-GFP (Addgene_48138, ^85^) or pSpCas9(BB)-2A-mCherrry (Addgene_197420, ^86^) vector. The successful insertion of the sgRNAs was verified by Sanger sequencing. Plasmids containing a sgRNA were transfected into HAP1 or HeLa cells with Lipofectamine 3000 (Thermo Fisher; Cat# L3000008). After 48 h, single GFP- and mCherry-positive cells were sorted by fluorescence-activated cell sorting (FACS) into 96 well plates (BD FACSMelody™ Cell Sorter). Single-cell colonies were expanded, and positive clones were identified by immunoblotting. Candidate knockout clones with loss of protein expression for the target of interest were further analyzed. After DNA extraction, the regions of interest surrounding the sgRNA target sequence were amplified by PCR and analyzed by Sanger sequencing. A detailed description is available (dx.doi.org/10.17504/protocols.io.81wgbw6kngpk/v1).

### Generation of PiggyBac cell lines

HeLa wild-type p40PX-EGFP (CVCL_F4CF) and ATG2 DKO p40PX-EGFP (CVCL_F4CG) cells, as well as HeLa wild-type p40PX-EGFP mCherry-LC3B (CVCL_F4CH) and ATG2 DKO p40PX-EGFP mCherry-LC3B (CVCL_F4CI) cells, were generated by co-transfecting cells with one or two PiggyBac (PB) transposon plasmids (Addgene_ 260697, Addgene_ 260698) together with a PB transposase expression vector. The PB transposase expression plasmid mPB (pBase) was obtained from the Wellcome Sanger Institute, UK under a material transfer agreement (MTA) and used as previously described in Cadiñanos and Bradley, 2007 (PMID: 17576687). HeLa cells were transfected using Lipofectamine™ 3000 (Thermo Fisher, Cat# L3000008) according to the manufacturer’s instructions. Following transfection, cells were selected using puromycin (Thermo Fisher, Cat# A1113803) and subsequently enriched by FACS sorting based on GFP or GFP/mCherry expression after treatment with 5 ng/mL doxycycline (Merck, Cat# D9891-5g). A detailed description is available (dx.doi.org/10.17504/protocols.io.x54v9qqnzl3e/v1).

### Protein expression and purification from *E. coli*

Recombinant proteins were expressed in E. coli Rosetta™ (DE3) pLysS cells (Novagen, Cat# 71403-3). Bacterial cultures were grown in 2x TY medium at 37 °C until reaching an optical density at 600 nm (OD600) of 0.4. The temperature was then reduced to 18 °C, and cultures were allowed to grow until an OD600 of 0.8 was reached. Protein expression was induced with 100 µM isopropyl β-D-1-thiogalactopyranoside (IPTG; Gerbu, Cat# 367-93-1) and allowed to proceed for 16 h at 18 °C. Cells were harvested by centrifugation, washed once with ice-cold 1x PBS, flash-frozen in liquid nitrogen, and stored at −80 °C until further use.

For the purification of glycine-exposed ATG8 proteins (Addgene_223728, Addgene_223729, Addgene_223730, Addgene_223726, Addgene_216836 and Addgene_223727), cell pellets were resuspended in lysis buffer containing 50 mM Tris-HCl pH 7.4, 300 mM NaCl, 1% Triton X-100, 5% glycerol, 2 mM MgCl₂, 1 mM DTT, 2 mM β-mercaptoethanol, 1x cOmplete™ EDTA-free protease inhibitor cocktail (Roche, Cat# 11836170001), Phosphatase Inhibitor Cocktail (Eubio, Cat# B15002), and DNase I (Sigma-Aldrich, Cat# DN25-1G). Cells were lysed by sonication, and lysates were clarified by centrifugation at 26000xg for 45 min at 4 °C using a Sorvall RC6+ centrifuge equipped with an F21S-8×50Y rotor (Thermo Scientific). The cleared supernatant was incubated with pre-equilibrated Glutathione Sepharose 4B beads (GE Healthcare, Cat# 17075605) for 2 h at 4 °C. Beads were washed twice with standard wash buffer (50 mM Tris-HCl pH 7.4, 300 mM NaCl, 1 mM DTT), once with high-salt wash buffer (same as above but with 700 mM NaCl), and twice again with standard buffer. Proteins were eluted overnight at 4 °C using 50 mM reduced glutathione (Sigma-Aldrich, Cat# G4251-50G) in wash buffer. After removal of beads by centrifugation and filtration (0.45 µM; Whatman, Cat# 6880-2504), the supernatant was concentrated using 10 kDa cut-off Amicon filters (Merck Millipore, Cat# UFC9010) and subjected to size-exclusion chromatography (SEC) using a Superdex 200 Increase 10/300 GL column (Cytiva, Cat# 28990944) equilibrated in SEC buffer (25 mM Tris-HCl pH 7.4, 300 mM NaCl, 1 mM DTT). Protein-containing fractions were identified by SDS-PAGE, pooled, concentrated, aliquoted, snap-frozen in liquid nitrogen, and stored at −80 °C. A detailed protocol is available (dx.doi.org/10.17504/protocols.io.kxygx4zdol8j/v1).

For the purification of GST- and mCherry-tagged glycine-exposed GABARAP (Addgene_244946), GST- and mCherry-tagged full-length GABARAP (Addgene_244947), GST-tagged full-length GABARAP (Addgene_244948), GST- and His-tagged glycine-exposed LC3B (Addgene_244950), and the GST-tagged LIR motif of p62 (amino acids 334– 342; Addgene_244939), cell pellets were resuspended in lysis buffer containing 50 mM HEPES pH 7.4, 300 mM NaCl, 1% Triton X-100, 5% glycerol, 2 mM MgCl₂, 1 mM DTT, 2 mM β-mercaptoethanol, 1x cOmplete™ EDTA-free protease inhibitor cocktail, Phosphatase Inhibitor Cocktail, and DNase I. Cells were lysed by sonication, and lysates were cleared by centrifugation at 38000xg for 45 min at 4 °C using a Sorvall RC6+ centrifuge (F21S-8×50Y rotor, Thermo Scientific). The cleared supernatant was incubated with pre-equilibrated Glutathione Sepharose 4B beads for 2 h at 4 °C. Beads were washed twice with standard wash buffer (50 mM HEPES pH 7.4, 300 mM NaCl, 1 mM DTT), once with high-salt wash buffer (same as above but with 700 mM NaCl), and twice again with standard buffer. For GST-tagged LIR motif (p62), the protein was eluted overnight at 4 °C using 50 mM reduced glutathione in wash buffer. For the other GST-tagged proteins, proteins were cleaved with TEV protease overnight at 4 °C in elution buffer. After removal of beads by centrifugation and filtration (0.45 µM), the supernatant was concentrated using 10 kDa cut-off Amicon filters and subjected to size-exclusion chromatography (SEC) using a Superdex 200 Increase 10/300 GL column equilibrated in SEC buffer (25 mM HEPES pH 7.4, 300 mM NaCl, 1 mM DTT). Protein-containing fractions were identified by SDS-PAGE, pooled, concentrated, aliquoted, snap-frozen in liquid nitrogen, and stored at −80 °C. A detailed protocol is available (dx.doi.org/10.17504/protocols.io.q26g7n4e3lwz/v1).

For purification of His-tagged mCherry-2xFYVE (Addgene_244944) cell pellets were resuspended in lysis buffer containing 50 mM HEPES (pH 7.4), 300 mM NaCl, 1% Triton X-100, 5% glycerol, 2 mM MgCl₂, 1 mM DTT, 2 mM β-mercaptoethanol, 1x cOmplete™ EDTA-free protease inhibitor cocktail, phosphatase inhibitor cocktail, and DNase I. Cells were lysed by sonication on ice, and lysates were clarified by centrifugation at 33000xg for 45 min at 4 °C using a Hitachi Himac CR22N centrifuge (R20A2 rotor). The supernatant was filtered and loaded onto a 5 mL HisTrap HP column (Cytiva), pre-equilibrated with water and Buffer A (30 mM HEPES pH 7.5, 300 mM NaCl, 10 mM imidazole, 2 mM β-mercaptoethanol). The column was washed with three column volumes of Buffer A, and bound proteins were eluted using a stepwise imidazole gradient (30, 75, 100, 150, 225, and 300 mM imidazole) by increasing the proportion of Buffer B (30 mM HEPES pH 7.5, 300 mM NaCl, 300 mM imidazole, 2 mM β-mercaptoethanol). Elution fractions were analyzed by SDS-PAGE and Coomassie staining. Fractions containing His-tagged mCherry-2xFYVE were pooled and concentrated using a 10 kDa molecular weight cut-off Amicon Ultra centrifugal filter, and further purified by size-exclusion chromatography on a Superdex 200 Increase 10/300 GL column equilibrated with SEC buffer (25 mM HEPES pH 7.5, 150 mM NaCl, 1 mM DTT). Protein-containing fractions were pooled, concentrated, aliquoted, snap-frozen in liquid nitrogen, and stored at –80 °C. A detailed protocol is available (dx.doi.org/10.17504/protocols.io.6qpvrw652lmk/v1).

A protocol for the purification of the E2-like enzyme ATG3 (Addgene_169079; (dx.doi.org/10.17504/protocols.io.btgknjuw) has been published previously.

### Protein expression and purification from insect cells

To purify His-tagged WIPI4 from insect cells (Addgene_244943) a codon-optimized synthetic gene was purchased from GenScript. The gene was cloned into a baculoviral expression vector, transformed to DH10EMBacY cells (Geneva Biotech) and the resulting bacmid DNA was verified by PCR for correct insertion of the transgene. Purified bacmid DNA was used to transfect *Spodoptera frugiperda* (Sf9) cells (Thermo Fisher Scientific, Cat# 12659017, RRID: CVCL_0549). For transfection, 2500 ng of bacmid DNA was mixed with FuGENE HD transfection reagent (Promega, Cat# E2311) and applied to 1 million Sf9 cells seeded in a 6-well plate. Seven days post-transfection, the first viral stock (V0) was harvested and used to infect 40 mL of Sf9 cells at 1×10⁶ cells/mL. Upon the first signs of cytopathic effect (viability drop and yellow fluorescence), the culture was centrifuged, and the supernatant containing the amplified virus (V1) was collected and stored at 4 °C. For large-scale expression, 1 L of Sf9 cells (1×10⁶ cells/mL) was infected with 1 mL of V1 virus. Cells were harvested when viability dropped to 90–95%, washed with PBS, flash-frozen in liquid nitrogen, and stored at – 80 °C until purification. Frozen pellets were thawed on ice and resuspended in 40 mL of lysis buffer (50 mM HEPES pH 7.5, 500 mM NaCl, 10% glycerol, 2 mM MgCl₂, 2 mM β-mercaptoethanol) supplemented with one cOmplete™ EDTA-free protease inhibitor tablet, 1 µL benzonase (Sigma-Aldrich, Cat# E1014-5KU), and a trace amount of DNase I per 50 mL. Cells were lysed by sonication (4 x 30 s at 60% amplitude with ice cooling). Lysates were clarified by centrifugation at 45000xg for 1 h at 4 °C (Beckman Coulter, TI45 rotor), and the supernatant was filtered through a 0.45 µM filter. Protein purification was performed using a 5 mL HisTrap HP column (Cytiva, Cat# 17524802), pre-equilibrated with water and Buffer A (30 mM HEPES pH 7.5, 300 mM NaCl, 10 mM imidazole, 2 mM β-mercaptoethanol). The column was washed with three column volumes of the same buffer, and His-tagged proteins were eluted using a stepwise imidazole gradient (30, 75, 100, 150, 225, and 300 mM) by increasing addition of Buffer B (30 mM HEPES pH 7.5, 300 mM NaCl, 300 mM imidazole, 2 mM β-mercaptoethanol). Elution fractions were analyzed by SDS-PAGE and Coomassie staining. Fractions containing His-tagged proteins were pooled and concentrated using a 10 kDa molecular weight cut-off Amicon Ultra filter, and further purified by size-exclusion chromatography using a Superdex 200 Increase 10/300 GL column equilibrated in SEC buffer (25 mM HEPES pH 7.5, 150 mM NaCl, 1 mM DTT). Protein-containing fractions were pooled, concentrated, aliquoted, snap-frozen in liquid nitrogen, and stored at –80 °C. A detailed protocol is available (dx.doi.org/10.17504/protocols.io.81wgbw6bqgpk/v1).

The ATG9A construct (Addgene_244940) was designed with an N-terminal 6xHis tag followed by a TEV cleavage site, the coding sequence for ATG9A, GFP, and a Twin-Strep tag. The ATG9A coding sequence was codon-optimized for insect cell expression and synthesized by GenScript. Purified bacmid DNA was used to transfect Sf9 cells (Thermo Fisher Scientific, Cat# 12659017, RRID: CVCL_0549). For transfection, 2500 ng of bacmid DNA was mixed with FuGENE HD transfection reagent and applied to 1 million Sf9 cells seeded in a 6-well plate. Ten days post-transfection, the first viral stock (V0) was harvested and used to infect 40 mL of Sf9 cells at 1×10⁶ cells/mL. Upon the first signs of cytopathic effect (viability drop and yellow fluorescence), the culture was centrifuged, and the supernatant containing the amplified virus (V1) and used to generate V2 virus by adding 1 mL of V1 to 30 mL of Sf9 cells at 1×10^6^ cells/mL. After three days 20 mL of V2 was used to infect 2 L of Sf9 cells (1×10^6^ cells/mL). Cells were harvested when viability dropped to 90–95% (usually five days after infection) washed with PBS, flash-frozen in liquid nitrogen, and stored at –80 °C until purification.

The cell pellets were resuspended by gentle pipetting and lysed using a Dounce homogenizer (10 strokes with loose pestle, followed by 2×20 strokes with tight pestle). The lysate was cleared by centrifugation at 7000xg (Sorvall RC6+ Centrifuge, F21S rotor) for 15 min. The cleared lysate was subjected to another round of centrifugation at 186000xg (Beckman XPN90, Ti45 rotor) for 1h at 4°C. The membrane containing pellet was resuspended in lysis buffer containing 0.3% DDM (DDM; Glycon, Cat# D97002-10g) using Dounce homogenizer and incubation at 4°C with gentle rolling. After 1h of incubation the insoluble material was removed by centrifugation at 186000xg (Beckman XPN90, Ti45 rotor) for 1h. The supernatant was loaded onto a 1 mL HisTrap HP column. The protein was eluted by a stepwise gradient of imidazole (40, 80, 150, 200 and 300 mM imidazole; 5 mL for each step) in 30 mM HEPES pH 7.5, 300 mM NaCl, 10% glycerol, 2 mM b-mercaptoethanol and 0,3 % DDM. Elution fractions were analyzed by SDS-PAGE and Coomassie staining. Fractions containing ATG9A-GFP were pooled and concentrated using a 100 kDa molecular weight cut-off Amicon Ultra Centrifugal filter (Merck Millipore, Cat# UFC8100), and further purified by size-exclusion chromatography using a Superose 6 Increase 10/300 GL column (GE Healthcare, Cat# 29091596) equilibrated in SEC buffer (25 mM HEPES pH 7.5, 150 mM NaCl, 1 mM DTT, 0.2% DDM). Protein-containing fractions were pooled, aliquoted, snap-frozen in liquid nitrogen, and stored at –80 °C. A detailed protocol is available (dx.doi.org/10.17504/protocols.io.yxmvmbn6bg3p/v1).

Protocols for the purification of the E1-like enzyme ATG7 (dx.doi.org/10.17504/protocols.io.bsennbde), the E3-like ligase complex ATG12– ATG5/ATG16L1β (Addgene_169076; dx.doi.org/10.17504/protocols.io.br6qm9dw), as well as untagged and mCherry-tagged PI3KC3-C1 (Addgene_187992, Addgene_187936; dx.doi.org/10.17504/protocols.io.8epv59mz4g1b/v1) have been published previously. The construct for ATG7 (pFastBac-mATG7) was a kind gift from Gammoh Noor.

### Protein expression and purification from HEK393F cells

To purify ATG2A and its variants from HEK293F cells, we expressed the protein from a pcDNA3.1(+) backbone encoding 3XFLAG-TEV-ATG2A (Addgene_244949) or 3XFLAG-TEV-ATG2A-mCherry (Addgene_244945). Mutations were introduced by *in vitro* mutagenesis to generate the ΔLIR mutants 3XFLAG-TEV-ATG2A-LIR-mCherry (Addgene_244951) and 3XFLAG-TEV-ATG2A-LIR-mCherry (Addgene_244952). The protein was expressed in FreeStyle^TM^ HEK293F cells (Thermo Fisher, Cat# R79007), grown at 37°C in FreeStyle^TM^ 293 Expression Medium (Thermo Fisher, Cat# 12338-026). The day before transfection, cells were seeded at a density of 0.7 x 10^6^ cells per mL. On the day of transfection, a 400 mL culture was transfected with 400 µg of the MAXI-prep DNA, diluted in 13 mL of Opti-MEM I Reduced Serum Medium (Thermo Fisher, Cat# 31985-062), and 800 µg Polyethylenimine PEI 25K (Polysciences, Cat# 23966-1), also diluted in 13 mL of Opti-MEM media. One day post transfection, the culture was supplemented with 100 mL EXCELL 293 Serum-Free Medium (Sigma-Aldrich, Cat# 14571C-1000ML). Another 72 h later, cells were harvested by centrifugation at 270 g for 20 min. The pellet was washed with PBS to remove medium and then flash-frozen in liquid nitrogen. For purification of ATG2A, pellets were thawed on ice in 20 mL of lysis buffer (50 mM HEPES pH 8.0, 300 mM NaCl, 10% glycerol, 1 mM TCEP, 0.5 M urea, 0.2% n-octyl-β-D-glucopyranoside (OG; Glycon, Cat# D97001-10g), supplemented with 1x cOmplete EDTA-free protease inhibitors (Roche), 1 µL Benzonase per 50 mL, and 200 µL Protease Inhibitor Cocktail per 50 mL (Sigma-Aldrich, Cat# P8849-5ML)) for 20 to 30 min. Pellets were resuspended by gentle pipetting and lysed using a Dounce homogenizer (10 strokes with loose pestle, followed by 2×20 strokes with tight pestle). Lysates were clarified by centrifugation at 18000xg for 30 min at 4 °C (HITACHI Himac CR22N, R20A2 rotor). Cleared supernatants were incubated with 0.6 mL pre-equilibrated anti-FLAG M2 affinity resin (Sigma-Aldrich, Cat# A2220-5ML) for 3 h at 4 °C with gentle mixing. The resin was washed three times with 13 mL of wash buffer containing glycerol (50 mM HEPES pH 8.0, 300 mM NaCl, 10% glycerol, 1 mM TCEP), followed by three washes with wash buffer lacking glycerol (same composition but without glycerol). Beads were then transferred to 1.5 mL tubes and incubated overnight at 4 °C with 200 µL of glycerol-free wash buffer containing 5 µL of TEV protease per tube. Alternatively, the protein can be eluted using 100 µL of FLAG peptide (Sigma-Aldrich, Cat# F3290-4MG) at a concentration of 4 mg/mL. The next day, eluates (Elution 1) were collected, and beads were washed once with 150 µL of wash buffer without glycerol to recover residual protein (Elution 2). Protein was aliquoted, snap-frozen in liquid nitrogen, and stored at –80 °C. All steps were performed at 4 °C or on ice to preserve protein integrity. OG was used from a 10% stock in water (CMC ≈ 0.6%) to facilitate solubilization of membrane-associated protein. A detailed protocol is available (dx.doi.org/10.17504/protocols.io.n92ld6zdng5b/v1).

### Purification of ATG9A-containing vesicles

Native ATG9A-positive compartments were isolated as previously described by Adriaenssens et al., 2025 ^53^. Briefly, cells from five 15 cm dishes were harvested, washed, and flash frozen. After thawing on ice, cells were lysed by repeated passage through a 26G needle in Vesicle Isolation Buffer (VIB; 20 mM HEPES pH 7.5, 150 mM NaCl, 250 mM sucrose, 1x cOmplete™ EDTA-free protease inhibitor cocktail, 20 mM β-glycerophosphate, 1 mM sodium orthovanadate, 1 mM sodium fluoride, and 1 mM EDTA pH 8.0). Following removal of nuclei and debris by low-speed centrifugation (2000 g, 6 min, 4 °C), the post-nuclear supernatant was incubated overnight with pre-equilibrated anti-FLAG beads (Sigma-Aldrich, Cat# A2220). Beads were then washed and vesicles were eluted using FLAG peptide (Sigma-Aldrich, Cat# F32920-4MG) in Elution Base Buffer (EBB; 20 mM HEPES pH 7.5, 150 mM NaCl, 1x cOmplete™ EDTA-free protease inhibitor cocktail, 20 mM β-glycerophosphate, 1 mM sodium orthovanadate, and 1 mM sodium fluoride). All steps were carried out on ice to preserve integrity of these structures. A detailed step-by-step protocol is available at dx.doi.org/10.17504/protocols.io.dm6gpzok8lzp/v1. For the lipidomic analysis, GFP-Trap beads (ChromoTek, Cat# GTA-20) were equilibrated with water and EBB by centrifugation at 4000xg for 2 min at 4 °C. Eluted ATG9A-positive vesicles were incubated with 40 µL of pre-equilibrated GFP-Trap beads for 2 h at 4 °C on a roller to allow binding. After incubation, the beads were stored overnight at 4 °C. The following day, beads were washed three times with 100 µL of 20 mM ammonium bicarbonate (pH 8.0; Sigma-Aldrich, Cat# 09830-500G), with each wash step performed by centrifugation at 4000xg for 3 min at 4 °C. After the final wash, beads were resuspended in 100 µL of 20 mM ammonium bicarbonate (pH 8.0) and stored at 4 °C until submission. For mass spectrometry analysis, magnetic GFP-Trap beads (20 µL slurry per sample; ChromoTek, Cat# GTMA-20) were acetylated to reduce peptide signals from co-digested matrix ligands, as described in dx.doi.org/10.17504/protocols.io.kxygxzexkv8j/v3. After acetylation, the beads were washed five times with 1 mL EBB. FLAG-eluted ATG9A-positive vesicles were then incubated with the acetylated, pre-washed GFP-Trap beads (20 µL per sample) overnight at 4 °C on a roller. The next day, beads were gently washed three times with 100 µL EBB. For submission, the beads were resuspended in 50 µL EBB. A detailed step-by-step protocol is available (PRIDE: PXD066668).

### Negative Stain Electron Microscopy

The isolated ATG9A compartments from cells grown in rich medium were applied on the carbon-coated copper-palladium grids (glow-discharged for 60 s immediately before use), stained twice for 30 s with 5 µL of 2 % uranyl acetate, air dried, and stored under vacuum until imaging. The stained grids were imaged using a FEI Morgagni 268D transmission electron microscope equipped with a tungsten filament emitter operated at 80 kV and an 11-megapixel Morada CCD camera (Olympus). The images were taken at 44 000x magnification and analysed with ImageJ (RRID:SCR_003070; https://imagej.net/). A detailed step-by-step protocol is available (dx.doi.org/10.17504/protocols.io.dm6gpmbn1gzp/v1).

### Dynamic Light Scattering

Dynamic light scattering (DLS) experiments were performed using a DynaPro NanoStar dynamic light scattering instrument (Wyatt Technology) using a quartz cuvette. Measurements were conducted in 20 mM HEPES pH 7.4, 150 mM NaCl and 1 mM TCEP). Samples included buffer alone, a MOCK sample (derived from HAP1 wild-type cells not expressing labeled ATG9A), or ATG9A-containing vesicles. All measurements were done at 25°C and each measurement consisted of 10 consecutive acquisitions of 5 seconds each. The scattering data were measured and analyzed with Dynamics 7.6.1.9 software (Wyatt Technology) using the globular protein model. A detailed step-by-step protocol is available (dx.doi.org/10.17504/protocols.io.yxmvmbnm5g3p/v1).

## Lipidomic analysis

ATG9A vesicles bound to GFP-Trap beads, as well as control beads subjected to mock purification using HAP1 wild-type cells, were prepared as described above. Whole-cell lysates (WCL) were generated from 1 x 10^6^ cells per sample. Cells were harvested by trypsinization and washed three times with PBS to remove residual media components, followed by three additional washes with 20 mM ammonium bicarbonate buffer (pH 8.0; Sigma-Aldrich, Cat# 09830) to eliminate PBS-derived salts that could interfere with downstream mass spectrometry. The final cell pellet was resuspended in 20 mM ammonium bicarbonate buffer and stored at 4 °C until submission. Vesicle pellets and pellets from WCL were resuspended in 260 µL methanol and spiked with 10 µL of lipid internal standard solution (SPLASH® Lipidomix®; Avanti, Cat# 330707) and deuterated C18:1 Glucosyl Ceramide-d5 standard (Avanti, Cat# 860673) and mixed by vortexing. Lipid fraction was extracted using a Matyash Modified Method: after addition of 1000 µL Methyl tert-butyl ether (MTBE) and vortexing, 200 µL water were added to induce phase separation. After centrifugation at 1000 g for 10 min, the upper organic phase was collected and dried under a soft nitrogen flow. Lipid extract was resuspended in 20 µL of methanol and transferred to vials with inserts for analysis. The LC-MS analysis was performed using a Vanquish UHPLC system (Thermo Fisher) combined with an Orbitrap Fusion™ Lumos™ Tribrid™ mass spectrometer (Thermo Fisher). Lipid separation was performed by reversed phase chromatography employing an Accucore C18, 2.6 µM, 150 x 2 mm (Thermo Fisher) analytical column at a column temperature of 35 °C. As mobile phase A an acetonitrile/water (50/50, v/v) solution containing 10 mM ammonium formate and 0.1 % formic acid was used. Mobile phase B consisted of acetonitrile/isopropanol/water (10/88/2, v/v/v) containing 10 mM ammonium formate and 0.1% formic acid. The flow rate was set to 400 µL/min. A gradient of mobile phase B was applied to ensure optimal separation of the analyzed lipid species. The mass spectrometer was operated in ESI-positive and -negative mode, capillary voltage 3500 V (positive) and 3000 V (negative), vaporization temperature 320 °C, ion transfer tube temperature 285 °C, sheath gas 60 arbitrary units (AU), auxiliary gas 20 AU and sweep gas 1 AU. Orbitrap MS scan mode at 120000 mass resolution was employed for lipid detection. The scan range was set to 250-1200 m/z for both positive and negative ionization mode, the AGC target was set to 2.0e5 and the intensity threshold to 5.0e3. The data analysis was performed using TraceFinder software (Thermo Fisher; RRID:SCR_023045). The raw data is publicly available via Zenodo.

### Lipidomics Data Analysis

Lipidomic data analysis was performed using Microsoft Excel (RRID:SCR_016137). Missing values in the raw data (reported in pmol) were replaced with zero. To account for background signals originating from the purification procedure, lipid abundances detected in the MOCK control samples were first averaged across the three biological replicates for each lipid species. The resulting mean MOCK values were then subtracted from the corresponding lipid abundances measured in each biological replicate of the ATG9A vesicle samples. Negative values resulting from background subtraction were set to zero. Following background correction, the total lipid content of each sample was calculated as the sum of all detected lipid species. The molar percentage (%mol) of each lipid species was then determined by dividing its background-corrected abundance by the total lipid content of the corresponding sample and multiplying by 100: %mol = (lipid species abundance / total lipid abundance) x 100. To obtain the final lipid composition, molar percentages were averaged across biological replicates for each lipid class. Lipid species were grouped into the following categories: Cholesterol, PC (including LPC and PC), PE (including LPE and PE), PI, PS, SM, and OTHERS (including CE, Cer, DAG, GlcCer, LacCer, PG, and TAG). A detailed protocol is available (dx.doi.org/10.17504/protocols.io.ewov11o27vr2/v1).

### Microscopy-based bead assay

Glutathione Sepharose 4B beads (GE Healthcare, Cat# 17075605) were used for binding GST-tagged bait proteins, and GFP-Trap agarose beads (ChromoTek, Cat# GTA-20) were used to bind GFP-tagged bait proteins.

For assays using ATG9A compartments or MOCK controls as prey (**Fig. 1E, 1G, 3A**), freshly purified ATG9A compartments (generated from 1 x 10^8^ cells) were incubated with 10 µL of pre-equilibrated GFP-Trap agarose beads. MOCK samples were prepared from HAP1 wild-type cells. Beads were equilibrated by washing once with water and twice with Elution Base Buffer (20 mM HEPES pH 7.5, 150 mM NaCl, 1x cOmplete™ EDTA-free protease inhibitor cocktail, 20 mM β-glycerophosphate, 1 mM sodium orthovanadate, and 1 mM NaF, in MilliQ H₂O, sterile-filtered and precooled). Binding was performed for 5 h at 4 °C with gentle rotation, and the samples were subsequently stored overnight at 4 °C. On the following day, beads were washed twice with 200 µL of SEC150 (25 mM HEPES pH 7.5, 150 mM NaCl, 1 mM DTT) and resuspended in 5 to 10 µL SEC150. For experiments (**Fig. 1G, S1H**) using ATG9A proteoliposomes (PLs) as bait, 500 µL of PLs were incubated with 10 µL of pre-equilibrated GFP-Trap agarose beads (washed with water and twice with SEC150) at 4 °C.

Glass-bottom 384-well microplates (Greiner Bio-One, Cat# 781892) were prepared with 20 µL of prey protein samples in SEC150. Beads (1 µL/well) were added and incubated with prey proteins for 30 min at room temperature (RT) prior to imaging. Prey proteins included mCherry-PI3KC3-C1 (0.1 µM) in **Fig. 1E and S1H** and ATG2A-mCherry (0.1 µM) in **Fig. 3A**. To assess PI3KC3-C1 activity (**Fig. 1G**), a cofactor mix was prepared (final concentrations: 0.5 mM ATP, 0.5 mM MgCl₂, 2 mM MnCl₂, 1 mM EGTA). ATG9A vesicle- or ATG9A PL-coated beads were first incubated with 0.03 µM PI3KC3-C1 for 30 min at RT, followed by 0.03 µM mCherry-2xFYVE for 10 min, before adding cofactors. The total reaction volume per well was 20 µL. For the bead-based lipid transfer assay (**Fig. 4B**), 125 µL of ATG9A PLs were incubated with 10 µM ATG2A and 6,25 µL of 0.5 mg/mL PI-containing donor SUVs for 2.5 hours at RT. After incubation, ATG9A PLs were bound to 4 µL of pre-equilibrated GFP-Trap agarose beads (slurry) for 2.5 h. Beads were washed once with SEC150, once with SEC250 (25 mM HEPES pH 7.5, 250 mM NaCl, 1 mM DTT), once with SEC350 (25 mM HEPES pH 7.5, 350 mM NaCl, 1 mM DTT) and twice with SEC150 to remove proteins and donor liposomes. The beads were treated with PI3KC3-C1, cofactors, and mCherry-2xFYVE as described above. To verify that the washing steps used in the assay shown in **Fig. 4B** are sufficient to efficiently remove ATG2A, 125 µL of ATG9A proteoliposomes (PLs) were incubated with 10 µM ATG2A-mCherry and 6.25 µL of 0.5 mg/mL PI-containing donor liposomes for 2.5 h at room temperature (**Fig. S5A**). After incubation, ATG9A PLs were immobilized on 4 µL of pre-equilibrated GFP-Trap agarose bead slurry for 2.5 h. Samples were collected before washing to determine the amount of ATG2A-mCherry bound to ATG9A PLs. Beads were then washed sequentially with SEC150, SEC250, SEC350, and twice with SEC150 to remove unbound ATG2A-mCherry. Additional samples were collected after washing to assess the fraction of ATG2A-mCherry that remained associated with the ATG9A PLs. To assess GABARAP lipidation on native ATG9A-containing vesicles (**Fig. S6F**), ATG9A compartments bound to GFP-Trap agarose beads were incubated for 2.5 h at 30 °C in 20 µL reactions containing 250 nM mCherry-GABARAPΔL117 (GABARAP-GLY), 250 nM ATG7 (E1), 250 nM ATG3 (E2), and 250 nM of a reconstituted E3 complex (comprising ATG5, ATG12, and ATG16L1), supplemented with 2 mM ATP and 1 mM MgCl₂. Reactions were performed in 0.5 mL tubes. After incubation, beads were washed once with SEC150. Negative controls included reactions using either mCherry-tagged full-length GABARAP (FL), which cannot be lipidated due to an uncleaved C-terminal residue, or mCherry alone, as well as reactions in which individual components of the ATG8 lipidation machinery (E1, E2, or E3) were omitted.

To examine the interaction between ATG2A and ATG8 (**Fig. 6A, S7A**), 5 µL Glutathione Sepharose 4B beads per condition were equilibrated with water and SEC150, then incubated with 5 µM GST-tagged bait protein in SEC150 for 1 h at 4 °C. After washing three times with SEC150, beads were resuspended in 5 µL SEC150. 384-well glass-bottom microplates were prepared with 20 µL samples containing 0.5 µM ATG2A-mCherry or ATG2A-LIR-mCherry, and 1 µL of beads was added per well.

To reconstitute lipid transfer to ATG9A-containing membranes in a microscopy-based bead assay, either ATG9A PL (**Fig. S5C**) or freshly purified ATG9A compartments (**Fig. 3E**) were used. For proteoliposomes, the lipid composition reflected that of isolated ATG9A compartments supplemented with a FRET pair (50% POPE, 20% Cholesterol, 26% DOPC, 1% Liver PI, 2% DOPS, 1% Sphingomyelin, 2% ATTO425-DOPE, 2% ATTO540Q-DOPE). For isolated ATG9A compartments, donor SUVs containing the FRET pair were used (66% DOPC, 30% DOPE, 2% ATTO425-DOPE, 2% ATTO540Q-DOPE). ATG9A PL or ATG9A compartments were incubated with 5 µL of GFP-Trap agarose beads pre-equilibrated in SEC150 buffer (25 mM HEPES pH 7.5, 150 mM NaCl, 1 mM DTT). After overnight binding with gentle rolling at 4 °C, beads were washed twice with 200 µL of SEC150 buffer and resuspended in 10 µL of the same buffer. Donor SUVs were prepared in SEC300 buffer (25 mM HEPES pH 7.5, 300 mM NaCl, 1 mM DTT), sonicated, extruded through a 100 nm filter, and stored at 4 °C overnight. For proteoliposome-based assays, donor SUVs lacked fluorophores (70% DOPC, 15% DOPE, 15% DOPS), whereas for compartment-based assays they contained the FRET pair as indicated above. For the assay, 1 µL of bead-bound material, 1.5 µL donor SUVs (0.5 mg/mL), and ATG2A (final concentration 250 nM) were added to wells of a glass-bottom 384-well microplate (Greiner Bio-One, Cat# 781892) in SEC300 buffer to a final reaction volume of 20 µL. Reactions were incubated in the dark at room temperature for 2 h prior to imaging. Detailed protocols for the transfer into ATG9A PL (dx.doi.org/10.17504/protocols.io.kqdg3mr5zl25/v1) and ATG9A compartments (dx.doi.org/10.17504/protocols.io.kxygxj64kl8j/v1) are available.

All samples were imaged using a Zeiss LSM 700 confocal microscope equipped with a Plan Apochromat 20x/0.8 WD 0.55 mm objective. A detailed protocol is available (dx.doi.org/10.17504/protocols.io.e6nvw4kr7lmk/v1).

### Conjugation of GABARAP to ATG9A compartments

Isolated ATG9A compartments and MOCK preparations were incubated with 10 µL GFP-Trap beads by mixing on a rotating wheel overnight at 4 °C. The beads were then washed twice with Reaction Buffer (25 mM HEPES, pH 7.5, 150 mM NaCl, 1 mM DTT). Each 20 µL reaction contained 2 mM ATP, 1 mM MgCl₂, and the following proteins at final concentrations: 250 nM hATG12–ATG5-ATG16L1, 250 nM mATG7, 250 nM hATG3, and 500 nM GABARAP. For lipidation experiments, we used a GABARAPΔL117 mutant, which lacks the C-terminal leucine and thus exposes the penultimate glycine required for PE conjugation. Full-length GABARAP, which cannot undergo lipidation, was included as a negative control. As a positive control, we used ATG9A PL containing 20.2% DOPC, 1.7% DOPS, 43.2% DOPE, 33.6% cholesterol, 1.3% sphingomyelin. Reactions were incubated at 30 °C for 3 h. After incubation, beads were washed once with 200 µL Reaction Buffer, resuspended in SDS loading buffer, and analysed by SDS-PAGE (15% gel). GABARAP and ATG9A were detected by Western blotting. A detailed protocol is available (dx.doi.org/10.17504/protocols.io.5qpvodbzzg4o/v1).

### Pull-Down Assay to Assess ATG2A Binding to WIPI4 and LC3B

FLAG trap beads (5 µl slurry per condition) were coated with purified 3xFLAG-tagged ATG2A by incubation in SEC300 buffer (25 mM HEPES pH 7.5, 300 mM NaCl, 1 mM TCEP). Beads were then incubated with 5 µM purified 6xHIS-tagged WIPI4, LC3B, or GFP (negative control) for 1 h at room temperature. After one wash with 100 µL SEC300, 40 µL SEC300 was added to the beads. Half of the sample was taken as the *low-salt* fraction. The remaining half was subjected to the following washing steps: two washes with SEC300, one wash with SEC500 (500 mM NaCl), and two final washes with SEC300. This sample was taken as the *high-salt* fraction. Samples were resuspended in SDS loading buffer and analyzed by SDS–PAGE. ATG2A (Proteintech, Cat# 23226-1-AP) and HIS-tagged proteins (Qiagen, Cat# 34660) were detected by Western blotting. A detailed protocol is available (dx.doi.org/10.17504/protocols.io.rm7vz9bw5gx1/v1).

### Analytical Size-Exclusion Chromatography of ATG2A-WIPI4 Complex

Analytical SEC was performed using a Superose 6 Increase 3.2/300 column (Cytiva, Cat# 29091598) pre-equilibrated with SEC buffer (25 mM HEPES pH 8.0, 300 mM NaCl, 1 mM TCEP). 3XFLAG-TEV-ATG2A and 6XHIS-WIPI4 were mixed at final concentrations of 4 µM and 4.8 µM, respectively, in a total volume of 50 µL. As controls, each protein was analyzed individually at the same concentrations and volume. The protein mixture was incubated on ice for 2 h before injection onto the column. Elution was monitored by absorbance at 280 nm, and 100 µL fractions were collected for subsequent Western blot analysis. ATG2A and WIPI4 were detected using anti-ATG2A (Proteintech, Cat# 23226-1-AP) and anti-6XHIS (Qiagen, Cat# 34660) antibodies. A detailed protocol is available (dx.doi.org/10.17504/protocols.io.ewov11oeovr2/v1).

### SDS-PAGE and western blot analysis

Samples were loaded on 4–12% SDS-PAGE gels (Invitrogen, Cat# NP0321BOX, or NP0323BOX) with PageRuler Prestained protein marker (Thermo Fisher, Cat# 26620, or 26617). Proteins were transferred onto nitrocellulose membranes (GE Healthcare, Cat# RPN132D) for 1.5 h using the Mini Trans-Blot Cell (Bio-Rad, Cat# 20-45-111). After the transfer, membranes were blocked with 5% milk powder dissolved in TBS-Tween (TBS-T, 0.1% Tween 20; Sigma-Aldrich, Cat# P7949-500mL) for 1 h at RT. The membranes were incubated overnight at 4°C with primary antibodies dissolved in the blocking buffer, washed three times for 5 min, and incubated with species-matched secondary horseradish peroxidase (HRP)-coupled antibodies diluted 1:10,000 in blocking buffer for 1 h at RT. Membranes were washed three times with TBS-T, incubated with SuperSignal West Femto Maximum Sensitivity Substrate (Thermo Fisher, Cat# 34096) and imaged with a ChemiDoc MP Imaging system (Bio-Rad). Images were analyzed with ImageJ (RRID:SCR_003070; https://imagej.net/). The primary antibodies used in this study are: anti-ATG2A (1:1000, Proteintech, Cat# 23226-1-AP), anti-ATG9A (1:1000, Cell Signaling, Cat# 5504; RRID:AB_2798241), anti-ATG14 (1:1000, Cell Signaling, Cat# 13509; RRID:AB_10695397), anti-BECLIN1 (1:1000, Cell Signaling, Cat# 3738; RRID:AB_490837), anti-CALNEXIN (1:1000, Transduction Laboratories, Cat# 610524; RRID:AB_397884), anti-GABARAP (1:1000, Cell Signaling Technology, Cat# 13733; RRID:AB_2798306), anti-GM130 (1:1000, BD Transduction Laboratories, Cat# 610823; RRID:AB_398142), anti-6XHIS (1:1000, Qiagen, Cat# 34660; RRID:AB_2619735), anti-LAMP1 (1:1000, Abcam, Cat# 25630; RRID:AB_470708), anti-LC3B (1:1000, Cell Signaling, Cat# 3868; RRID:AB_2137707 or NanoTools, Cat# 0260-100/LC3-2G6; RRID:AB 2943418), anti-p62/SQSTM1 (1:1000, BD, Cat# 610833; RRID:AB_398152), anti-RAB7 (1:1000, Cell Signaling Technology, Cat# 9367; RRID:AB_1904103), anti-TOM20 (1:1000, Santa Cruz Biotechnology, Cat# sc-17764; RRID:AB_628381), anti-TGN46 (1:1000, Novus Biologicals, Cat# H00010618-M02; RRID:AB_830312). The secondary antibodies used in this study are: HRP conjugated polyclonal goat anti-mouse (Jackson ImmunoResearch Labs, Cat# 115-035-003; RRID:AB_10015289) and HRP conjugated polyclonal goat anti-rabbit (Jackson ImmunoResearch Labs, Cat# 111-035-003; RRID:AB_2313567). Uncropped and unprocessed scans of blots are provided in the Source Data file. A detailed protocol is available (dx.doi.org/10.17504/protocols.io.n2bvje64wgk5/v1).

### Immunofluorescence and confocal microscopy

Cells were seeded on glass coverslips (12 mm #1.5; Marienfeld, Cat# 0117520) at a density of 100,000 cells per well (6 well plate). The following day, cells were transiently transfected using Lipofectamine™ 3000 (Thermo Fisher, Cat# L3000008) according to the manufacturer’s instructions (2500 ng DNA). For transient transfection we used DNA for EGFP-p40PX (kind gift from Michael Yaffe, Addgene_19010), EGFP-2xFYVE (Addgene_244941), ATG2A wild-type (Addgene_244949) and EGFP-WIPI4 (Addgene_223769). Where indicated in the figure legends, 24 h post-transfection, cells were treated for 2 h with either Earle’s balanced salt medium (EBSS; Sigma-Aldrich, Cat# E3024-500ML) and 100 nM Bafilomycin A1 (BafA; Santa Cruz Biotechnology, Cat# sc-201550), or 2 µM VPS34-IN1 (ApexBio, Cat# APE-B6179). Cells were then fixed in cold methanol for 5 min at –20 °C. For EEA1 staining, fixation was performed using 4% paraformaldehyde for 15 min at room temperature. After fixation, cells were washed with PBS and then permeabilized and blocked simultaneously for 60 min at RT in a solution of PBS containing 0.25% (v/v) Triton™ X-100 (Sigma-Aldrich, Cat# T8787) and 5% (v/v) Bovine Serum Albumin (BSA; Sigma-Aldrich, Cat# A9647).

Primary and secondary antibodies were diluted in the same blocking buffer and incubated for 2 h each at RT, with three PBS washes between each step. Coverslips were mounted onto microscopy slides using DAPI Fluoromount-G® (Southern Biotech, Cat# 0100-20), which counterstains nuclei, and stored at –20 °C until imaging. Confocal microscopy was performed with a Zeiss LSM900 equipped with an Airyscan 2 module and a Plan-Apochromat 63x/1.4 Oil DIC, WD 0.19 mm objective. The images were acquired and processed with 2D Airyscan processing plug-in in Zen Blue software (Zeiss). The primary antibodies used in this study are: anti-ATG9A (1:100, Cell Signaling Technology, Cat# 13509; RRID:AB_2798241), anti-ATG14 (1:100, Cell Signaling Technology, Cat# 5504; RRID:AB_10695397), anti-ATG16L1 (1:100, Cell Signaling Technology, Cat# 8089; RRID:AB_10950320), anti-EEA1 (1:100, Cell Signaling Technology, Cat# 3288; RRID:AB_2096811), anti-EGFP (1:100, ChromoTek, Cat# 3h9; RRID:AB_10773374), anti-LC3B (1:100, NanoTools, Cat# 0260-100/LC3-2G6; RRID:AB_2943418), anti-p62/SQSTM1 (1:100, Abnova Cat# H00008878-M01; RRID:AB_437085), anti-ULK1 (1:100, Cell Signaling Technology Cat# 8054; RRID:AB_11178668) and anti-WIPI2 (1:100, Bio-Rad Cat# MCA5780GA; RRID:AB_10845951). The secondary antibodies used in this study are: AlexaFluor-488 goat anti-rat IgG (H+L) (1:500, Thermo Fisher Scientific Cat# A-11008, RRID: AB_2338047), AlexaFluor-546 goat anti-mouse IgG (H+L) (1:500, Thermo Fisher, Cat# A-11003; RRID: AB_2534071), AlexaFluor-647 goat anti-rabbit IgG (H+L) (1:500, Thermo Fisher, Cat# A-21244; RRID: AB_2535812). A detailed protocol is available (dx.doi.org/10.17504/protocols.io.4r3l21okxg1y/v1).

### Live-cell imaging

Cells were seeded in 10-well glass-bottom imaging chambers (Greiner Bio-One, Cat# 543079) at 6000 cells per well and treated with 5 ng/mL doxycycline (Merck, Cat# D9891-5g). The following day cells were treated with 2 µM VPS34-IN1 inhibitor (ApexBio, Cat #APE-B6179) or EBSS, and imaged using a Visitron Live Spinning Disk microscope (CFI Plan Apo Lambda 100x/1.45 Oil, WD 0.13 mm objective, sCMOS camera, 488 nm laser diode (150 mW) and 561 nm solid state laser (200 mW); including a temperature & CO2-controlled environment). A detailed protocol is available (dx.doi.org/10.17504/protocols.io.ewov1re8plr2/v1).

### Generation of liposomes and proteoliposomes

Small unilamellar vesicles (SUVs) for tethering, lipid transfer assays, and ATG9A reconstitution were prepared using lipid compositions as indicated in the figure legends. The chloroform-dissolved lipids were mixed in a glass vial, dried under an argon stream, and further desiccated under vacuum for 1 h. The resulting lipid film was rehydrated in either SEC300 buffer (25 mM HEPES pH 7.5, 300 mM NaCl, 1 mM TCEP) for tethering and lipid transfer assays or SEC150 buffer (25 mM HEPES pH 7.5, 150 mM NaCl, 1 mM DTT) for ATG9A proteoliposome (PL) preparation. Rehydrated lipids were gently mixed and sonicated for 2 min in a bath sonicator, followed by extrusion through a 0.1 µM membrane (Whatman, Cat# 10419504) using a Mini Extruder (Avanti, Cat# 610024). The resulting SUVs were used at a final lipid concentration of 1 mg/mL. The lipids used in this study are: ATTO425-DOPE (ATTO-TEC, Cat# AD 425-161), ATTO540Q-DOPE (ATTO-TEC, Cat# AD 540Q-161), DOPC (Avanti, Cat# 850375C), DOPE (Avanti, Cat# 850725C), DOPS (Avanti, Cat# 840035C), POPE (Avanti, Cat# 850757C-25MG), PI3P (Sigma-Aldrich, Cat# 850150P-500UG), Liver PI (Sigma-Aldrich, Cat# 840042C-25MG), NBD-PE (Sigma-Aldrich, 810155P-1MG), Rh-PE (Invitrogen, Cat# L-1392), DGS-NTA (Avanti, Cat# 790404C), Cholesterol (Avanti, Cat# 700000P-100mg), Sphingomyelin (Avanti, Cat# 860584P-5MG) and ATTO390-PE (ATTO-TEC, Cat# AD 390-161). The final concentration of the SUV suspension was 0.5 mg lipids/mL. The compositions of SUVs used in our experiments are provided in the figure legends, where percentages are indicated as a shorthand for mol%. For ATG9A incorporation, SUV suspensions were adjusted to 2% CHAPS (Glycon, Cat# D99009-25G) and incubated for 1 h at RT. ATG9A-GFP, purified in micelles containing 0.2% DDM, was added at a final concentration of 0.5 µM (1:1 volume ratio with the SUV-CHAPS mixture) and incubated for 1 h at RT. The mixture was then diluted with SEC150 to reduce detergent concentration below the critical micelle concentration (final DDM concentration of ∼0.003% and final CHAPS concentration of 0.07%). Proteoliposome reconstitution was completed by overnight dialysis at 4 °C against SEC150 containing 0.1 g/l Bio-Beads SM-2 (Bio-Rad, Cat# 1523920), followed by a 1 h incubation with fresh Bio-Beads at RT. Non-incorporated material was removed by centrifugation at maximum speed in a benchtop centrifuge for 30 min. The supernatant containing ATG9A PLs was collected and used for downstream experiments. A detailed protocol is available (dx.doi.org/10.17504/protocols.io.x54v95y6ml3e/v1).

### Tethering and lipid transfer assay

Lipid transfer assays were performed at room temperature in glass-bottom 384-well microplates. Each reaction contained donor and acceptor liposomes (2.5 µL each; 0.5 mg/mL total lipids) and the indicated proteins (250 nM ATG2A, 100 nM 6xHis-WIPI4, and 100 nM 6xHis-LC3B). 6xHis-WIPI4 was recruited to SUVs via its PI3P-binding domain, whereas 6xHis-LC3B was recruited through DGS-NTA lipids incorporated into the SUVs. Reactions were carried out in SEC300 buffer in a final volume of 50 µL. Fluorescence was recorded every 20 seconds for 1.5 h using a Tecan SPARK Multimode Microplate Reader (Tecan Life Sciences; RRID:SCR_021897), with excitation at 485 nm and emission at 535 nm for the NBD–Rhodamine B FRET pair, or with excitation at 405 nm and emission at 470 nm for the ATTO425–ATTO540Q FRET pair. At the end of each reaction, 0.5% (v/v) DDM was added to fully solubilize lipids and determine the maximal fluorescence, measured 50 min after DDM addition. Fluorescence values were normalized by subtracting the initial fluorescence and dividing by the maximum fluorescence after DDM addition.

In parallel, liposome tethering was monitored in the same wells by measuring absorbance at 405 nm every 20 seconds over the course of the reaction using the same plate reader. This allowed quantification of vesicle clustering under identical buffer and experimental conditions. A detailed protocol is available (dx.doi.org/10.17504/protocols.io.eq2ly4nkplx9/v1).

### Isolation of Autophagosomes and Proteinase K Protection Assay

For the isolation of native autophagosomes (APs) or precursor structures, HAP1 ATG9A-GFP (RRID:CVCL_E2TR) and HAP1 ATG9A-GFP ATG2 DKO (CVCL_F0LE) cells were cultured in 15 cm dishes (two per condition). Cells were treated with EBSS and 100 nM BafA for two hours and harvested by trypsinization, pelleted by centrifugation, and washed with PBS. Cell pellets were then resuspended in 800 µL Vesicle Isolation Buffer (VIB; 20 mM HEPES pH 7.5, 150 mM NaCl, 250 mM sucrose, 1x cOmplete™ EDTA-free protease inhibitors, 20 mM β-glycerophosphate, 1 mM sodium orthovanadate, 1 mM NaF, and 1 mM EDTA pH 8.0) and transferred to LoBind tubes (Eppendorf, Cat# 0030108116). Cells were lysed by passing the suspension 30x through a 26G needle, followed by chilling on ice for 10 min, and an additional 30x passage. Lysates were cleared by centrifugation at 1,000xg for 10 min at 4 °C to pellet nuclei and cell debris. The supernatant was collected, and the pellet was re-extracted with 700 µL VIB, followed by a second 1,000 xg spin. Supernatants from both spins were pooled and centrifuged at 2,000xg for 5 min to remove additional debris, and the resulting supernatant was centrifuged at 10,000xg for 20 min at 4 °C to pellet the autophagosome-enriched fraction. The pellet was washed once with 500 µL wash buffer (WB) containing 20 mM HEPES pH 7.5 and 150 mM NaCl (with 1x cOmplete EDTA-free protease inhibitors), and re-pelleted under the same conditions. Finally, the pellet was resuspended in 500 µL WB (with 1 mM DTT) and used for downstream applications. To assess protease accessibility of isolated APs, a proteinase K protection assay was performed. 150 µL of autophagosome suspension were incubated with 10 μg/mL proteinase K (Thermo Fisher, Cat# EO0491) for 20 min at 4 °C. Digestion was stopped by the addition of 1 mM PMSF (Santa Cruz Biotechnology, Cat# sc-482875) and incubation on ice for 10 min. APs were collected by centrifugation at 10,000xg for 10 min at 4 °C, resuspended in 15 µL WB and resuspended in 6x SDS loading buffer, and analyzed by western blotting. A detailed protocol is available (dx.doi.org/10.17504/protocols.io.bp2l6z1q1gqe/v1).

### AlphaFold 3

We employed AlphaFold 3 ^87^ to predict potential interactions between ATG2A (Q2TAZ0-1), LC3B (H3BTL1), and WIPI4 (Q9Y484-1). Pairwise predictions were run using five models per prediction. Predictions with an inter-protein predicted TM-score (ipTM) > 0.5 were considered indicative of potential interactions. For each prediction, predicted aligned error (PAE) plots, pLDDT scores, and the resulting 3D structural models were manually inspected to assess confidence and binding interfaces. Structural models were visualized using UCSF ChimeraX (1.10). A detailed protocol is available (dx.doi.org/10.17504/protocols.io.6qpvr8rm2lmk/v1).

### Mass spectrometry data comparison and Venn diagram analysis

To assess protein overlap between datasets, the top 1,000 proteins from two replicates of mass spectrometry analysis were compared. Data processing and comparison were performed using Microsoft Excel. The COUNTIF function was used to determine overlaps between datasets, and the number of shared proteins was used to construct a Venn diagram to visualize common and unique protein identifications across the two datasets. A detailed protocol is available (dx.doi.org/10.17504/protocols.io.14egnr24ql5d/v1).

### Quantification of microscopy-based bead assay

Each experimental condition was analyzed in at least three independent biological replicates. Signal quantification from microscopy images was performed using an artificial intelligence (AI)-based analysis pipeline. The script automatically identifies individual beads and extracts signal intensities by drawing line profiles across bead surfaces, calculating the difference between the minimum and maximum gray values along each line. Bead segmentation was performed using Cellpose (RRID:SCR_022332)^88^, which had been trained specifically to recognize beads in our imaging setup. The source code is available at https://www.maxperutzlabs.ac.at/research/facilities/biooptics-light-microscopy. A detailed protocol is available (dx.doi.org/10.17504/protocols.io.rm7vz9b6rgx1/v1). Background-corrected intensity values were plotted and used for statistical analysis. For experiments with low signal-to-noise ratios (indicated in the figure legends), manual quantification was performed in ImageJ (Fiji). A detailed protocol is available (dx.doi.org/10.17504/protocols.io.x54v95d7zl3e/v1).

### Quantification of lipid transfer to ATG9A compartments

Signal quantification from microscopy images of three independent experiments (**Fig. 3E, 3F**) was performed using an artificial intelligence (AI)-based analysis pipeline as described above (“Quantification of microscopy-based bead assay”). Intensities in the red (ATTO540Q) and blue (ATTO425) channels were quantified to monitor SUV tethering and lipid transfer, respectively. The blue signal was corrected for bleed-through from ATG9A-GFP bound to the beads. The corrected blue signal reflects unquenched and de-quenched ATTO425, which increases upon lipid transfer due to separation from the quencher. The red signal (ATTO540Q), corresponding to the quencher present in donor SUVs, serves as a measure of SUV tethering to ATG9A compartments. To assess lipid transfer, the ratio of ATTO425 (blue, transfer) to ATTO540Q (red, tethered SUVs) was calculated, thereby normalizing transfer to the amount of bound donor SUVs. The resulting values were further normalized to the condition without ATG2A, in which no lipid transfer is expected, and residual blue signal arises from incompletely quenched ATTO425 in donor SUVs. Thus, normalized values above 1 indicate dequenching of ATTO425 due to increased distance from ATTO540Q, consistent with ATG2A-mediated lipid transfer. Values from three independent experiments were averaged, plotted, and used for statistical analysis. A detailed protocol is available (dx.doi.org/10.17504/protocols.io.kxygxj64kl8j/v1).

### Quantification of lipid transfer to ATG9A PL using ATTO425/ATT0540Q FRET pair

Signal quantification from microscopy images of three independent experiments (**Fig. S5C**) was performed using an artificial intelligence (AI)-based analysis pipeline as described above (“Quantification of microscopy-based bead assay”). The fluorescence signal measured at 405 nm excitation was normalized to the ATG9 PL-only control, which was set to 1 and is indicated by the dotted line in the graph (**Fig. S5D**). In this condition, the signal derives from two components: bleed-through of GFP in ATG9A-GFP and a residual ATTO425 signal being incompletely quenched. These signal components are present in all tested conditions, as the ATTO425/ATTO540Q FRET pair is included in the lipid composition of ATG9A PL. Values above 1 therefore indicate additional dequenching of ATTO425 due to increased separation from ATTO540Q, consistent with ATG2A-mediated lipid transfer. Values from three independent experiments were averaged, plotted, and used for statistical analysis. A detailed protocol is available (dx.doi.org/10.17504/protocols.io.kqdg3mr5zl25/v1).

### Co-localization Analysis

Co-localization analysis of immunofluorescence (IF) images was performed using the Pearson’s correlation coefficient (PCC) to assess signal overlap between fluorescent channels. Image analysis was carried out in Fiji (ImageJ) using the Just Another Co-localization Plugin (JaCoP) ^89^. PCC values were calculated for each image to quantify the linear correlation between the intensities of the two channels across all pixels. A PCC value close to +1 indicates strong positive co-localization, whereas values near 0 reflect random distribution. At least 2 images per condition per replicate were analyzed, and results were plotted and statistically evaluated using PRISM software (GraphPad Software). A detailed protocol is available (dx.doi.org/10.17504/protocols.io.bp2l6z98zgqe/v1).

## Data availability

The raw data generated in this study is publicly available on Zenodo (https://doi.org/10.5281/zenodo.20068186). The mass spectrometry proteomics data have been deposited to the ProteomeXchange Consortium via the PRIDE ^90^ partner repository with the dataset identifier PXD066668. Plasmids generated and used in this study are available from Addgene. A comprehensive overview of data, protocols, and key laboratory materials − including their persistent identifiers − is provided in the Key Resources Table, available as a supplementary file via Zenodo. Source data are provided as a Source Data file. All other data supporting the findings of this study are available from the corresponding author upon reasonable request. For the purpose of open access, the authors have applied a CC-BY 4.0 public copyright license to all Author Accepted Manuscripts arising from this submission.

## Code availability

No code was generated for this study.

## Acknowledgements

We thank members of the Martens lab and colleagues of the Aligning Science Across Parkinson’s (ASAP) Mito911 Team for their help and advice. We thank the Max Perutz Labs BioOptics, Flow Cytometry, and Mass Spectrometry facilities for technical support. Proteomics analyses were performed by the Mass Spectrometry Facility at Max Perutz Labs using the VBCF instrument pool. We thank the Vienna BioCenter Core Facilities (VBCF) Protech Facility for help with HEK cell expressions. We thank the Proteomics team of the Molecular Discovery Platform at CeMM for mass spectrometric data acquisition. We thank Viktoria Blahova for help with ATG2A-ATG8 interaction experiments, Stephen Shoebridge for support with the statistical analysis, and Luca Ferrari for support with EM of ATG9A compartments.

An earlier version of this manuscript was posted to bioRxiv on 2025-08-18 at https://doi.org/10.1101/2025.08.16.670665.

## Author Contributions Statement

E.H., J.S.M., and S.M. conceived the project. E.H., J.S.M., D.B., J.R., and S.M. designed the experiments. E.H., J.S.M., D.B., M.S., and J.R. performed the experiments. E.H., J.S.M., and S.M. wrote the original draft, and all authors contributed to editing and reviewing the manuscript.

---

**Supplementary Fig. 1.**
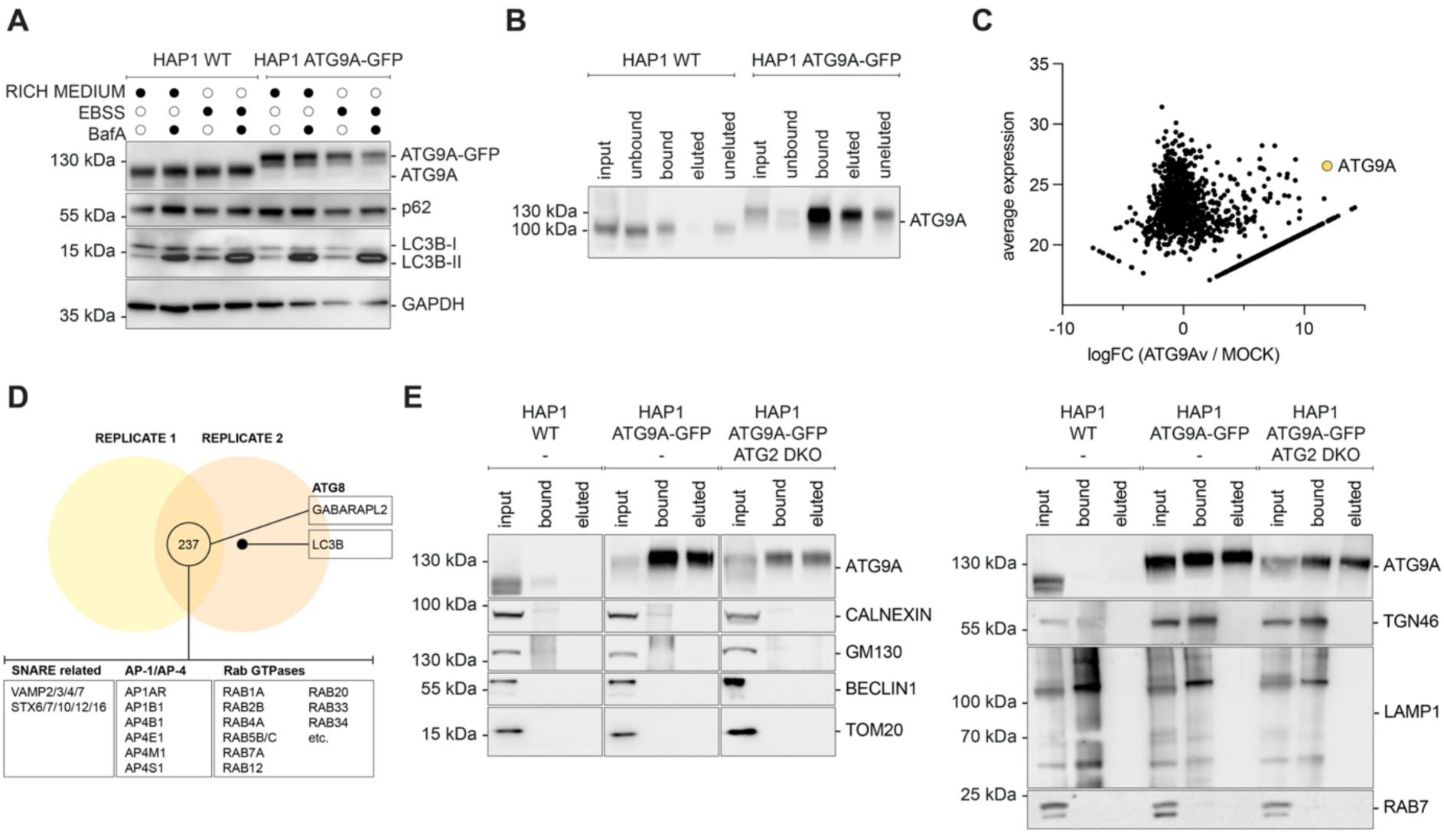
Isolation and analysis of native ATG9A compartments. (**A**) Western blot analysis of cell lysates from HAP1 wild-type and HAP1 ATG9A-GFP knock-in cell lines grown in rich medium or EBSS, either untreated or treated with 100 nM Bafilomycin A1 (BafA1) for 2 hours. Both the wild-type and knock-in cells display comparable levels of autophagy flux (LC3B), indicating that the GFP and FLAG tags do not impair ATG9A functionality. (**B**) Western blot analysis of fractions collected during the isolation of native ATG9A compartments. ATG9A is strongly enriched in the final eluate from the HAP1 ATG9A-GFP knock-in cell line, while it is nearly undetectable in the corresponding HAP1 wild-type (WT) control, indicating successful and specific enrichment of ATG9A compartments. (**C**) Volcano plot showing mass spectrometry results. ATG9A (yellow) is significantly enriched in vesicles isolated from HAP1 ATG9A-GFP cells compared to HAP1 wild-type (MOCK) cells, confirming the specificity of the isolation procedure. (**D**) Venn diagram showing the overlap of proteins identified in two independent replicates of ATG9A compartment isolations, highlighting shared proteins among the top 1000 hits. (**E**) Western blot analysis of input (0.5% of sample loaded), bound (2.8% of sample loaded), and eluted fractions (2.8% of sample loaded) from ATG9A vesicle isolations performed using HAP1 wild-type, ATG9A-GFP, and ATG9A-GFP ATG2 DKO cells. Approximately 1.2 × 10⁸ cells were lysed per isolation. ATG9A is enriched in the bound fraction from ATG9A-GFP and ATG9A-GFP ATG2 DKO cells, whereas markers of other membrane compartments, including EEA1, GM130, BECLIN1, TOM20, TGN46, LAMP1, and RAB7, are not co-purified.

**Supplementary Fig. 2.**
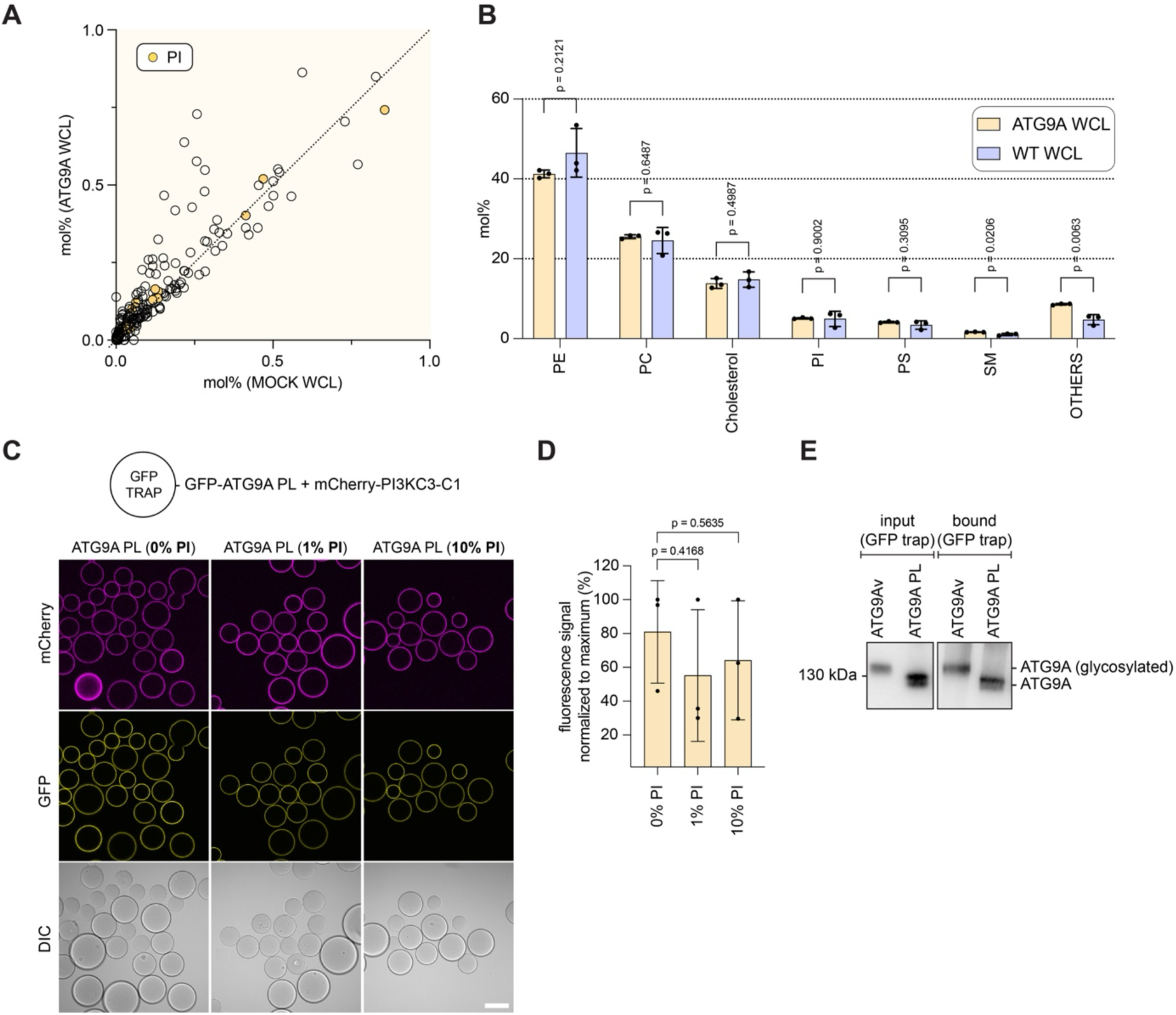
Analysis of native ATG9A compartments. (**A**) Scatter plot comparing the molar percentage (mol %) of individual lipid species in whole cell lysates of HAP1 wild-type cells (x-axis) and ATG9A-GFP cells (y-axis). Each point represents a lipid species. The clustering along the diagonal indicates a high similarity in lipid composition between the two cell lines. PI species are highlighted in yellow. (**B**) Lipidomic analysis of whole cell lysates (WCL) from HAP1 ATG9A-GFP (yellow) and wild-type (violet) cells. Lipid species were grouped by class, and their molar percentages were summed to yield the total mol% per lipid class shown on the x-axis. CE, Cer, DAG, GlcCer, LacCer, PG, and TAG were summarized under "others". Statistical significance within each class was assessed using an unpaired *t*-test. Bars represent mean ± s.d. (**C**) GFP-Trap beads coated with ATG9A PL were incubated with mCherry-PI3KC3-C1 (30 nM) to assess whether recruitment of PI3KC3-C1 depends on PI. **(D)** Quantification of mCherry fluorescence from panel **C** (*N* = 3; independent experiments). Statistical significance was assessed using an unpaired *t*-test. Bars represent mean ± s.d. mCherry signal (background-corrected) was expressed relative to the maximum value of each replicate. **(E)** Comparison of ATG9A-GFP levels in native ATG9A-containing vesicles and reconstituted ATG9A PLs. ATG9A-containing vesicles were isolated from HAP1 ATG9A-GFP cells. Reconstituted ATG9A PL were composed of 50% POPE, 20% cholesterol, 27% DOPC, 2% DOPS, and 1% sphingomyelin. For each condition, 300 µL of sample were subjected to GFP-Trap pull-down using 4 µL GFP-Trap bead slurry. Bound fractions were analyzed by Western blot. The higher apparent molecular weight of ATG9A-GFP in vesicle-derived samples is due to its glycosylation in mammalian cells, whereas the lower apparent molecular weight observed in PL samples reflects ATG9A purified from insect cells.

**Supplementary Fig. 3.**
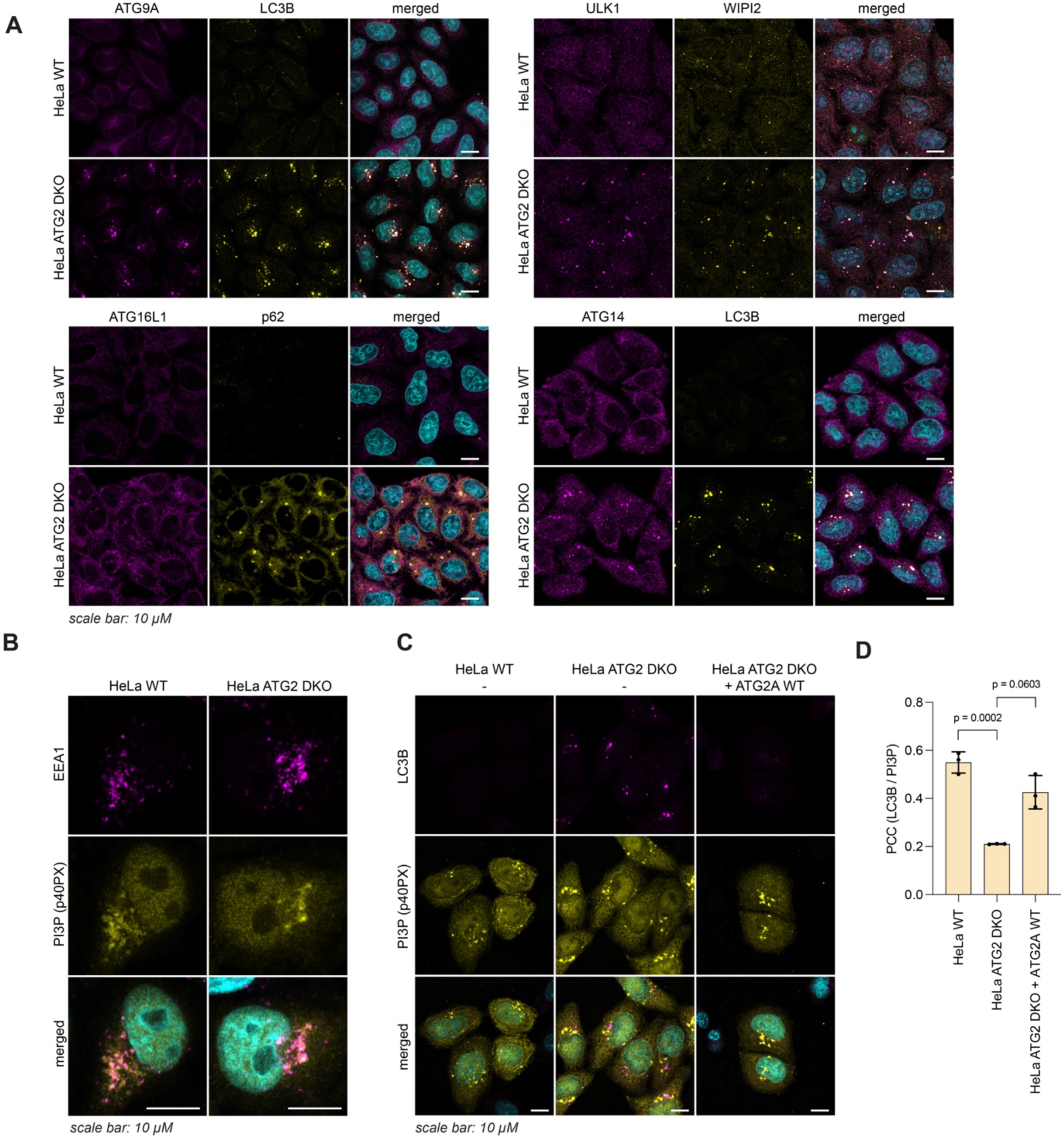
Foci formation in HeLa ATG2 DKO cells. (**A**) Immunofluorescence staining of HeLa wild-type and ATG2 DKO cells for autophagy-related proteins: ATG9A, LC3B, ATG16L1, p62, ULK1, WIPI2, and ATG14. ATG2 DKO cells show aberrant accumulation of autophagy markers in distinct foci. (**B**) Immunofluorescence staining of HeLa wild-type and ATG2 DKO p40PX-EGFP (PI3P-binding domain) for analyzing the co-localization of EEA1 and p40PX as a control for specificity of the detector (**C**) Rescue of foci phenotype in ATG2 DKO cells 24 h after transient transfection with a wild-type ATG2A construct. LC3B localization resembles that of HeLa wild-type cells, indicating restoration of normal trafficking. **(D)** Quantification of co-localization between LC3B and p40PX-EGFP in HeLa WT, ATG2 DKO cells, and ATG2 DKO cells transfected with a wild-type ATG2A construct (shown in **C**), assessed using Pearson’s correlation coefficient (PCC). Statistical significance was determined using an unpaired *t*-test (*N =* 3; independent experiments; 3 images per condition and replicate). Bars represent mean ± s.d.

**Supplementary Fig. 4.**
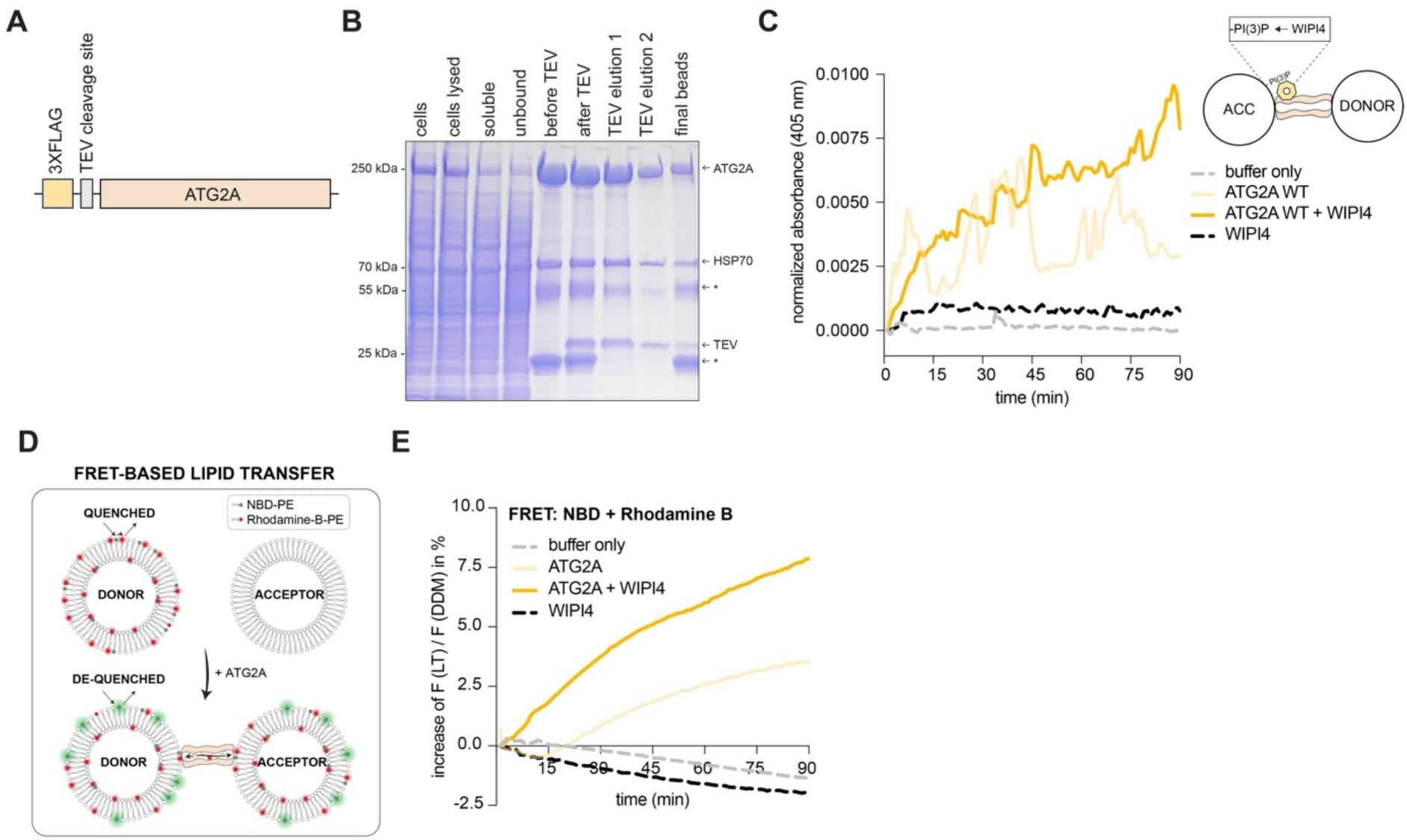
Purification of ATG2A. (A) Schematic of expression construct used to produce ATG2A and ATG2A ΔLIR in HEK293T cells. A separate construct containing a C-terminal mCherry tag was employed in bead-based microscopy assays. (**B**) Coomassie-stained SDS-PAGE showing protein fractions from purification workflow. The fractions labeled “TEV elution 1” and “TEV elution 2” contain the cleaved and purified ATG2A protein. A common co-purifying contaminant, HSP70, and the TEV protease, are indicated on the gel. The two other bands derive from the anti-FLAG beads (**C**) Liposome tethering assay measured by absorbance at 405 nm. Lines represent mean values (*N =* 3; independent experiments). Conditions include: no protein (buffer only), 250 nM ATG2A wild-type (WT), and 250 nM ATG2A wild-type in presence of 100 nM WIPI4 or 100 nM WIPI4 alone. Data reflect increased turbidity upon liposome clustering. (**D**) Schematic of FRET-based lipid transfer assay using donor and acceptor SUVs. NBD fluorescence increases upon dilution of NBD-PE into acceptor membranes, indicating lipid transfer. (**E**) Comparison of lipid transfer activity between ATG2A WT (250 nM) in absence or presence of WIPI4 (100 nM), or WIPI4 alone (100 nM). NBD fluorescence (F(LT)) was recorded at 535 nm (excitation at 485 nm) and normalized to fluorescence after membrane solubilization with 0.5% DDM (F(DDM)). Acceptor SUVs contained 65% DOPC, 15% DOPS, 15% DOPE, and 5% PI3P whereas donor SUVs contained 66% DOPC, 30% DOPE, 2% NBD-DOPE, and 2% Rhodamine B-DOPE. All SUVs were extruded through a 100 nm filter. Experiments were performed in triplicate; lines represent mean values (*N =* 3; independent experiments).

**Supplementary Fig. 5.**
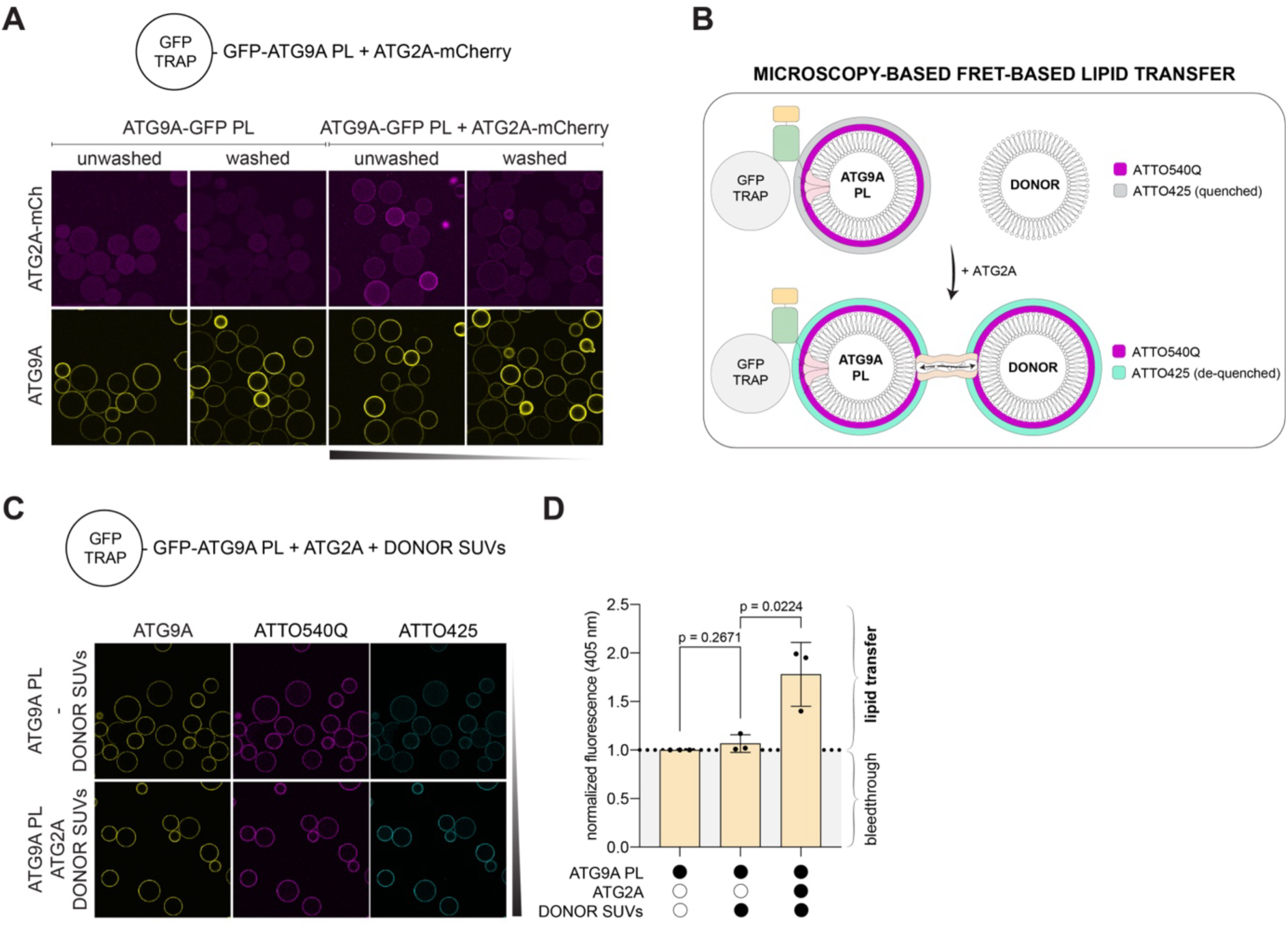
Lipid transfer and tethering assays with ATG2A. (**A**) ATG9A PL were incubated with ATG2A-mCherry for 2.5 hours, incubated with GFP-Trap agarose beads for 2.5 hours for immobilization of ATG9A PL and washed with high salt buffer. ATG9A PLs were composed of 50% POPE, 20% Cholesterol, 27% DOPC, 0% Liver PI, 2% DOPS, 1% Sphingomyelin (100 nm in size). (**B**) Schematic of the microscopy-based FRET-based lipid transfer assay using ATG9A PL including a ATTO425/ATTO540Q FRET pair and donor SUVs. ATTO425 fluorescence increases upon dilution of ATTO425-PE into donor membranes, indicating lipid transfer. (**C**) GFP-Trap agarose beads coated with ATG9A PL were incubated with donor SUVs (70% DOPC, 15% DOPE, 15% DOPS; 100 nm in size) in the presence or absence of ATG2A. The lipid composition of ATG9A PL reflected that of isolated ATG9A compartments supplemented with a ATTO425/ATTO540Q FRET pair (19% DOPC, 54% DOPE, 2% DOPS, 19% cholesterol, 1% sphingomyelin, 1% liver PI, 2% ATTO425-DOPE, 2% ATTO540Q-DOPE; 100 nm in size). (**D**) Quantification (*N =* 3; independent experiments) of ATTO425 fluorescence signal of the lipid transfer experiment shown in **C**. The fluorescence signal was normalized to the ATG9 PL-only control, which was set to 1 and is indicated by the dotted line. In this condition, the signal derived from the bleed-through of GFP containing ATG9A-GFP. Values above the line reflect ATTO425 dequenching caused by increased separation from ATTO540Q, resulting from ATG2A-mediated lipid transfer. Statistical significance was assessed using an unpaired *t*-test. Bars represent mean ± s.d.

**Supplementary Fig. 6.**
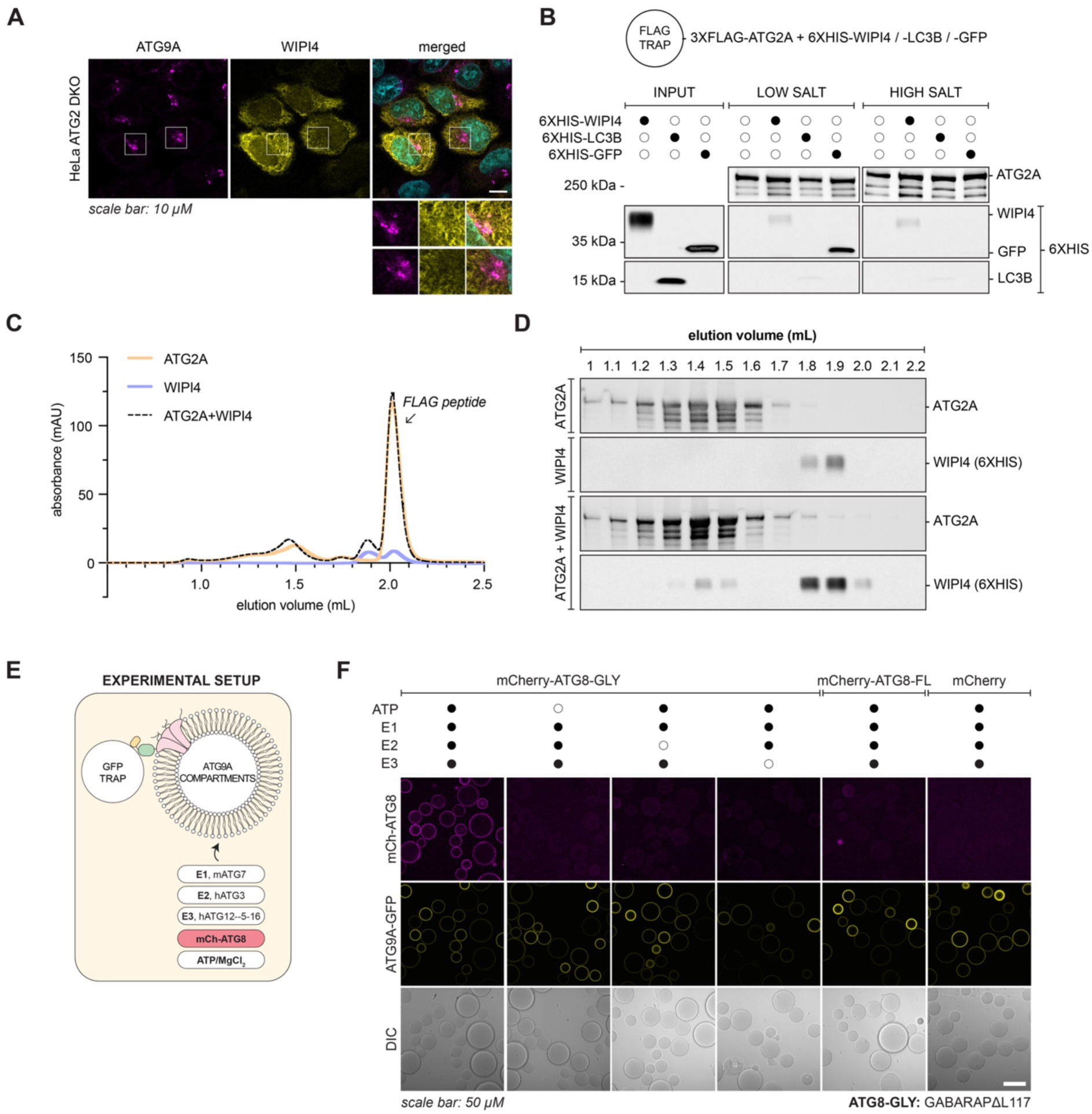
ATG9A compartments can serve as substrates for ATG8 lipidation. (A) Immunofluorescence staining of HeLa ATG2 DKO cells 24 h after transient transfection with EGFP-WIPI4. An antibody against EGFP was used to visualize WIPI4. Compared to other autophagy proteins (**Fig. S3A**), WIPI4 shows minimal localization to ATG9A-positive foci in ATG2 DKO cells, indicating that little or no PI3P is present at these sites. (**B**) FLAG Trap beads coated with 3×FLAG-ATG2A were incubated with 6×HIS-tagged WIPI4, LC3B, or GFP (negative control). After one wash with buffer containing 300 mM NaCl, signals were detected for all proteins (low-salt condition). Following additional washes, the GFP signal was lost, whereas WIPI4 and LC3B signals persisted (high-salt condition). Compared to the input, signals of bound proteins were relatively weak. WIPI4, GFP, and LC3B were detected using an anti-HIS antibody. (**C**) Superose 6 size-exclusion chromatography (SEC) profiles of ATG2A, WIPI4, or a mixture of ATG2A (4 µM) and WIPI4 (4.8 µM). (**D**) Western blot analysis of SEC fractions from ATG2A, WIPI4, or ATG2A-WIPI4 mixtures. WIPI4 shows partial co-elution with the ATG2A peak, but most remains unbound. (**E**) Schematic of the experimental setup shown in **F**. (**F**) GFP-Trap agarose beads coated with native ATG9A-GFP compartments were incubated with mCherry-GABARAPΔL117 (GABARAP-GLY) in the presence of the ATG8 lipidation machinery: E1 (ATG7), E2 (ATG3), and E3 (a complex of ATG12–ATG5-ATG16L1), along with ATP, to assess lipidation. mCherry fluorescence on the bead surface indicates successful conjugation of GABARAP to ATG9A compartments. As negative controls, beads were incubated with either mCherry-GABARAP full-length (FL), which cannot be lipidated, or mCherry alone.

**Supplementary Fig. 7.**
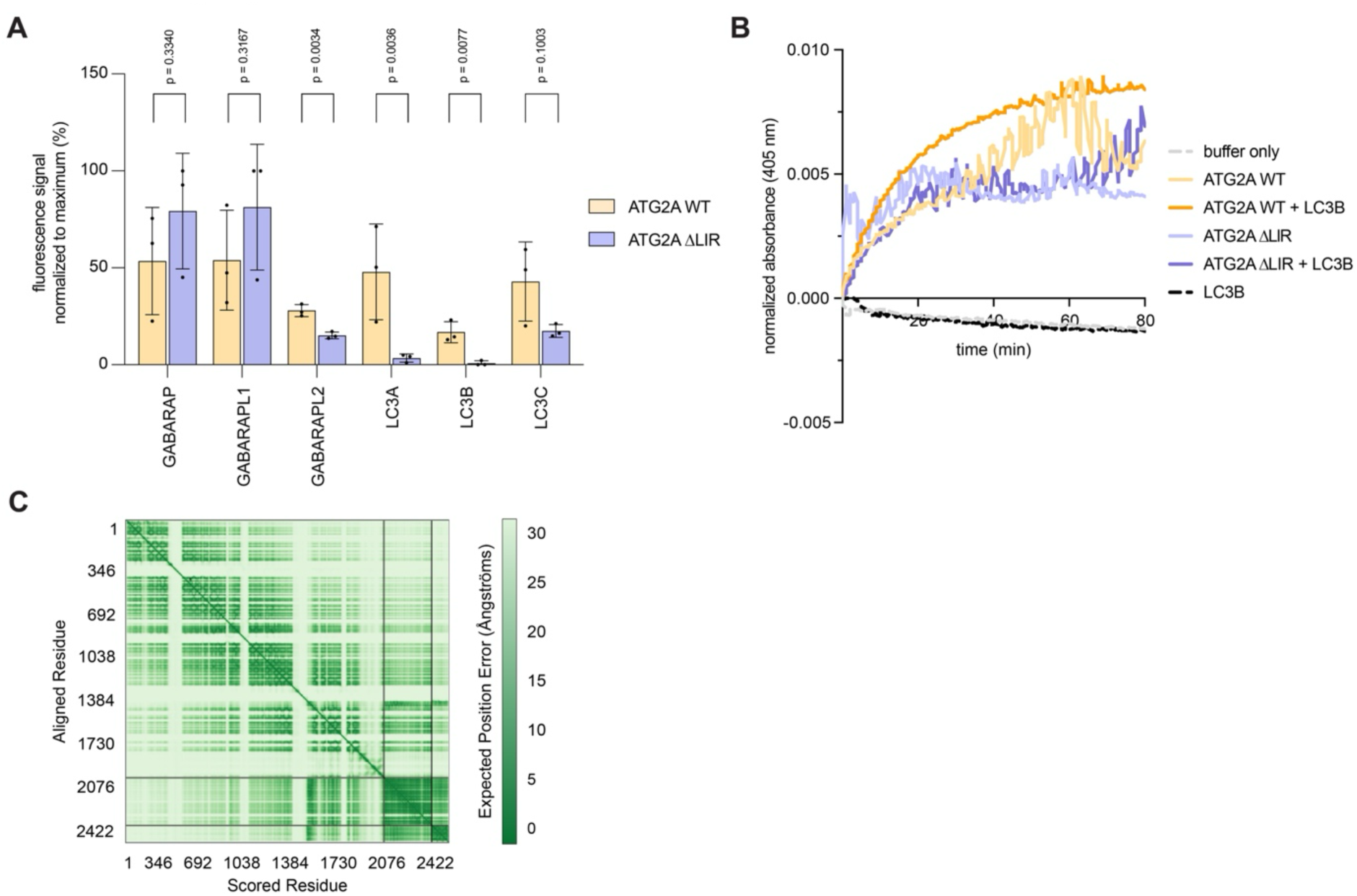
ATG8 directly interacts with ATG2A and enhances its lipid transfer activity. (**A**) Quantification of mCherry fluorescence signal from a microscopy-based bead assay assessing the recruitment of ATG2A to ATG8 proteins (as shown for LC3B in Fig. 6A). GSH-Trap beads coated with GST-tagged ATG8 family members (GABARAP, GABARAPL1, GABARAPL2, LC3A, LC3B, and LC3C) were incubated with either mCherry-tagged ATG2A or ATG2A-ΔLIR. Statistical significance was assessed using an unpaired *t*-test. Total fluorescence signals were normalized within each replicate to the respective maximum value. Each dot represents the mean of images acquired per replicate (*N =* 3; independent experiments). Bars represent mean ± s.d. (**B**) Liposome tethering assay measured by absorbance at 405 nm. Lines represent mean values (*N =* 4; independent experiments). Conditions include: no protein (buffer only), 250 nM ATG2A wild-type (WT) or mutant (ΔLIR), and 250 nM ATG2A wild-type or mutant (ΔLIR) in the presence of 100 nM 6XHIS-LC3B or 100 nM 6XHIS-LC3B alone. Data reflect increased turbidity upon liposome clustering, indicative of tethering activity (**C**) The predicted alignment error (PAE) heatmap for the AlphaFold3-predicted structure of the ATG2A-WIPI4-LC3B complex shown in Fig. 6D.

